# Reciprocal 16p11.2 Copy Number Variants Differentially Alter Regional Brain Anatomy and Allometric Scaling

**DOI:** 10.64898/2026.11.24.768986

**Authors:** S Prem, RT Shinohara, J Seidlitz, AZ Tomlinson, M Gardner, L Dorfschmidt, S Karandikar, D Zimmerman, KY Sun, TP Roberts, A Raznahan, B Jung, A Alexander-Bloch

## Abstract

Reciprocal 16p11.2 copy-number variants (CNVs) confer substantial risk for neuropsychiatric disorders and produce opposing effects on brain size, with larger brain volume in deletion carriers and smaller volume in duplication carriers. Regional alterations broadly parallel this global phenotype, but because regional brain measures scale nonlinearly with brain size, it remains unclear whether regional differences reflect expected consequences of altered brain size, regional anatomy disproportionate to brain size, or fundamental alterations in allometric scaling. We analyzed 285 imaging sessions from 235 participants (49 deletion, 59 duplications, 127 controls) using log-log allometric models to distinguish these possibilities. Brain-size adjustment revealed widespread cortical disproportion in both CNVs, with predominantly reciprocal effects driven by surface area rather than cortical thickness, whereas subcortical structures showed convergent reductions disproportionate to brain size. Strikingly, examining genotype-specific scaling exponents revealed distinct allometric patterns between the CNVs. In duplication, regional differences predominantly reflected fixed shifts relative to brain size. In deletion, by contrast, regional differences more frequently varied with brain size, with substantial exponent deviations affecting 32% of disproportionate cortical regions compared with 12% in duplication. Thus, reciprocal 16p11.2 dosage does not simply shift regional anatomy in opposite directions: duplication largely preserves the normally conserved scaling of regional anatomy with brain size, whereas deletion more extensively disrupts it.

## Introduction

Copy number variants (CNVs) at the proximal 16p11.2 (BP4–BP5) locus are among the most penetrant genetic risk factors for psychiatric and neurodevelopmental disorders, conferring increased risk for conditions including autism, intellectual disability, and schizophrenia ^(1–5)^. A striking feature of these reciprocal variants is their mirrored effect on brain size, with larger total brain volume in deletion carriers and smaller volume in duplication carriers ^(6–8)^. Neuroimaging studies have also identified widespread, broadly reciprocal cortical and subcortical alterations in both CNVs ^(8–10)^. However, it is unclear whether these regional differences are expected consequences of altered global brain size or are disproportionate to brain size.

Prior studies have addressed the influence of brain size using proportional normalization or linear regression to adjust regional measures for total intracranial volume (TIV), total surface area (SA), or mean cortical thickness (CT) ^(8–12)^. These approaches identified regional differences that persist after adjustment, suggesting that 16p11.2 CNVs (16p CNVs) have effects beyond their global brain-size phenotype. However, brain regions have established nonlinear scaling relationships with overall brain size ^(13–16)^, such that regional volumes do not necessarily increase in direct proportion to TIV ^(17–21)^. Allometric adjustment accounts for these relationships when determining whether regional differences are disproportionate to brain size ^(22–30)^. For example, studies of sex chromosome aneuploidies have used normative allometric scaling relationships to distinguish regional anatomical differences attributable to altered brain size from disproportionate changes, revealing selective subcortical and cerebellar abnormalities ^(28, 29)^.

While allometric adjustment accounts for nonlinear relationships between regional anatomy and overall brain size, it typically assumes that scaling relationships are shared across groups. This assumption is consistent with the remarkable conservation of regional brain-size scaling in normative populations ^(15, 16, 31, 32)^. Such conservation suggests that regional scaling relationships are not arbitrary but may instead reflect fundamental constraints on brain growth and organization arising from neurodevelopment ^(13, 33–37)^. Given that 16p CNVs disrupt early neurodevelopmental processes such as proliferation, migration, and differentiation ^(38–43)^, they may also alter scaling relationships, making allometric scaling itself a potentially informative neurodevelopmental phenotype. Scaling alterations could take two forms: a change in the scaling constant (intercept), producing a fixed regional shift at a reference brain size, or a change in the scaling exponent (slope), a more fundamental departure from normative scaling in which regional differences vary with brain size. Despite their reciprocal effects on overall brain size, whether deletion and duplication produce corresponding or distinct alterations in allometric scaling remains unknown.

Here, using structural MRI data from 16p CNV carriers and controls in the Simons Variation in Individuals Project (Simons VIP), we first used allometric adjustment to identify regional differences disproportionate to brain size and characterize the regional pattern of deletion and duplication effects. We further examined SA and CT, which jointly contribute to cortical volume but have distinct developmental and genetic determinants, to characterize the anatomical basis of these effects ^(44–46)^. Second, we tested whether 16p CNVs altered scaling exponents and compared the aggregate magnitude of scaling-exponent deviation between deletion and duplication. Finally, we integrated these analyses to determine how scaling-exponent deviations affected the interpretation of regional disproportion and whether this differed between deletion and duplication.

## Methods

### Participants

Participants were drawn from the Simons VIP (now Simons Searchlight), as previously described ^(10, 47)^. The sample included carriers of the recurrent 16p11.2 BP4–BP5 deletion or duplication and familial and unrelated noncarrier controls. The final sample comprised 235 participants (49 deletion carriers, 59 duplication carriers, and 127 controls, including 28 familial controls) contributing 285 imaging sessions (Tables 1–2). The study was approved by the institutional review boards of participating sites, and informed consent was obtained from participants or their legal representatives. Inclusion criteria and analytic sample preparation are described in the Supplementary Methods.

**Table 1:** Sample and imaging characteristics. Abbreviations: CTRL, control; DEL, 16p11.2 deletion; DUP, 16p11.2 duplication; W, White; B, Black; AAPI, Asian American and Pacific Islander; O, Other. Values are mean (SD) unless otherwise indicated. Age is at each participant’s first scan; statistical comparisons are participant-level. Race categories were not mutually exclusive. Other includes American Indian/Alaska Native, other, and more than one race. Only significant Tukey post-hoc comparisons are shown.

| **Measure** | **Control** | **16p11.2 Deletion** | **16p11.2 Duplication** | **Comparison** | **Statistic** | ***p*** | **Post-hoc tests** |
|---|---|---|---|---|---|---|---|
| Sessions | 127 | 67 | 91 | Descriptive only | - | - | - |
| Subjects | 127 | 49 | 59 | Descriptive only | - | - | - |
| Families | 120 | 41 | 34 | Descriptive only | - | - | - |
| Race data available, n (%) | 99 (78%) | 46 (94%) | 34 (58%) | Descriptive only | - | - | - |
| Race, n (% available) | W: 64 (65%)<br>B: 31 (31%)<br>AAPI: 6 (6%)<br>O: 14 (14%) | W: 41 (89%)<br>B: 2 (4%)<br>AAPI: 2 (4%)<br>O: 7 (15%) | W: 34 (100%)<br>B: 2 (6%)<br>AAPI: 0 (0%)<br>O: 2 (6%) | Descriptive only | - | - | - |
| Age range, years | 3.2–64.1 | 2.4–48.7 | 2.1–63.8 | Descriptive only | - | - | - |
| Female n (%) | 55 (43%) | 20 (41%) | 24 (41%) | CTRL vs DEL vs DUP | χ²(2) = 0.16 | 0.924 | - |
| Age, years | 24.2 (15.2) | 14.4 (10.2) | 27.1 (16.6) | CTRL vs DEL vs DUP | F(2, 232) = 11.29 | <0.001 | CTRL-DEL, p<0.001;<br>DEL-DUP, p<0.001 |

**Table 2.** Clinical and cognitive characteristics. Abbreviations: FSIQ, full-scale IQ; NVIQ, nonverbal IQ; VIQ, verbal IQ; ADOS CSS, Autism Diagnostic Observation Schedule calibrated severity score; ASD, autism spectrum disorder; BMI, body mass index. Values are mean (SD) or n/N with available data (%). ADOS ≥ 4 and clinical ASD diagnosis were compared with Fisher’s exact test. Head circumference Z was compared between DEL and DUP only. Only significant Tukey post-hoc comparisons are shown.

| **Measure** | **Control** | **16p11.2 Deletion** | **16p11.2 Duplication** | **Comparison** | **Statistic** | ***p*** | **Post-hoc tests** |
|---|---|---|---|---|---|---|---|
| N | 127 | 49 | 59 | Descriptive only | - | - | - |
| FSIQ | 107.7 (13.9) | 86.9 (13.4) | 89.4 (19.3) | CTRL vs DEL vs DUP | F(2, 215) = 43.40 | <0.001 | CTRL-DEL, p<0.001;<br>CTRL-DUP, p<0.001 |
| NVIQ | 105.2 (13.2) | 91.2 (12.4) | 89.0 (19.5) | CTRL vs DEL vs DUP | F(2, 216) = 28.29 | <0.001 | CTRL-DEL, p<0.001;<br>CTRL-DUP, p<0.001 |
| VIQ | 107.9 (14.3) | 83.4 (16.9) | 92.1 (18.4) | CTRL vs DEL vs DUP | F(2, 214) = 44.96 | <0.001 | CTRL-DEL, p<0.001;<br>CTRL-DUP, p<0.001;<br>DEL-DUP, p=0.017 |
| ADOS CSS | 1.3 (0.6) | 3.1 (2.2) | 2.4 (1.8) | CTRL vs DEL vs DUP | F(2, 130) = 8.27 | <0.001 | CTRL-DEL, p<0.001;<br>CTRL-DUP, p=0.024 |
| ADOS ≥ 4 | 0/27 (0%) | 14/49 (29%) | 10/57 (18%) | CTRL vs DEL vs DUP | Fisher's exact | 0.004 | - |
| Clinical ASD dx | 0/28 (0%) | 9/49 (18%) | 6/59 (10%) | CTRL vs DEL vs DUP | Fisher's exact | 0.035 | - |
| Head circumference Z | - | 1.8 (1.3) | -0.3 (1.4) | DEL vs DUP | Welch t(99.3) = 7.49 | <0.001 | - |
| BMI Z | 0.6 (1.0) | 1.4 (1.1) | 0.5 (1.0) | CTRL vs DEL vs DUP | F(2, 221) = 13.54 | <0.001 | CTRL-DEL, p<0.001;<br>DEL-DUP, p<0.001 |
| Height Z | 0.2 (1.1) | 0.3 (1.0) | 0.1 (1.2) | CTRL vs DEL vs DUP | F(2, 224) = 0.67 | 0.511 | - |
| Weight Z | 0.6 (1.0) | 1.4 (1.3) | 0.5 (1.2) | CTRL vs DEL vs DUP | F(2, 224) = 10.81 | <0.001 | CTRL-DEL, p<0.001;<br>DEL-DUP, p<0.001 |

### MRI acquisition

T1-weighted MRI data were obtained using research multi-echo or clinical single-echo MPRAGE protocols across participating sites ^(47)^. Detailed acquisition parameters and scanner information are provided in the Supplementary Methods.

### Image processing, quality control, and harmonization

T1-weighted images were processed using the FreeSurfer recon-all-clinical pipeline (v7.4.1), which is agnostic to scan orientation, contrast, and resolution ^(48–52)^. Cortical grey matter volume (cGMV), surface area (SA), and cortical thickness (CT) were extracted for 68 Desikan–Killiany parcels ^(53)^; non-cortical volumes were derived from the FreeSurfer aseg parcellation ^(54)^ using the integrated SynthSeg+ pipeline ^(48, 49)^. Global measures included TIV, cortical (cGMV) and subcortical grey matter volume (sGMV), white matter volume (WMV), ventricular (Vent) and cerebrospinal fluid (CSF) volume, corpus callosum volume (CC), mean CT, and total cortical SA.

Imaging-derived phenotypes were harmonized across scanners using Longitudinal ComBat, with age, sex, and genotype as fixed effects and participant as a random effect, preserving biological variation of interest and within-participant correlation ^(55)^.

Scans with a SynthSeg+ QC score below 0.65 or Euler characteristic below −60 were excluded before harmonization ^(49, 56, 57)^. Mean bilateral Euler number was included as an image-quality covariate in all analyses. QC and raw neuroanatomical distributions by genotype are shown in Figures S1-S2.

### Statistical analysis

Unless otherwise specified, analyses used linear mixed-effects models with nested random intercepts for participant within family ^(58, 59)^. Genotype was modeled as a three-level factor with controls as the reference, yielding deletion–control and duplication–control contrasts. Models included a natural spline for age (3 df), sex, and mean bilateral Euler number. Effect sizes were approximated as Cohen’s 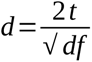. Statistical significance was defined as Benjamini–Hochberg false discovery rate (FDR)-adjusted *p* < 0.05, with analysis-specific correction families described in Supplementary Methods ^(61)^. Analyses were performed in R version 4.5.2 ^(62)^; figures were generated using ggplot2 version 4.0.1 and ggseg version 1.6.8 ^(63, 64)^.

### Disproportionate volume differences

We first estimated absolute regional genotype effects without TIV adjustment, then identified disproportionate effects by modeling log-transformed regional volume as a function of genotype and log-transformed TIV. This log-log formulation models the allometric relationship between regional volume and TIV while estimating genotype effects under a common slope ^(16, 22, 23, 27–29)^. Sensitivity analyses compared log-log adjustment with linear TIV adjustment and proportional normalization (volume/TIV, standardized by sample SD; Fig. S3).

To classify joint deletion and duplication effects, log-adjusted regional effects were grouped as canonical mirror (deletion increase, duplication decrease), reverse mirror (deletion decrease, duplication increase), convergent (same-direction), deletion-specific, duplication-specific, or no effect. Effects with *|d|* ≤ 0.10 were considered negligible; genotype-specific patterns required a non-negligible effect in one genotype and a negligible effect in the other. Classification was independent of statistical significance and applied to all 68 cortical parcels and 29 aseg structures with summaries focused on the 21 subcortical structures excluding ventricular and cerebral WMV.

Spatial correspondence between deletion and duplication effect-size maps was assessed using Spearman correlations. For cortical maps, significance was evaluated using two-sided spherical rotation tests as described in Supplementary Methods ^(65, 66)^.

### Surface area and cortical thickness

We repeated the disproportion analysis for SA and CT using total SA and mean CT as their respective global references. To check this choice, we compared, in controls, how much of each cortical region’s variance was explained by its own global measure versus TIV (Supplementary Methods; Fig. S4). Regional SA followed total SA and TIV similarly (median partial R² 0.32 vs 0.26), whereas regional CT followed mean CT but was essentially unrelated to TIV (0.16 vs 0.004). Consistent with this, SA effects were nearly unchanged with TIV as the reference, whereas CT effects were attenuated (Fig. S4B, C).

### Altered brain-size scaling

To test whether 16p genotype altered the normative scaling assumption, we compared three nested linear mixed-effects models, successively allowing the scaling constant and scaling exponent to vary by genotype:

### Common

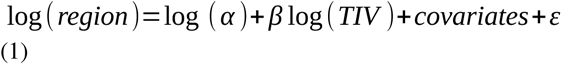

### Genotype-specific constant

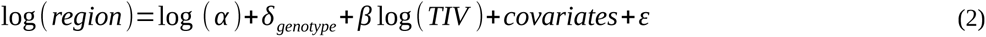

### Genotype-specific constant and scaling exponent

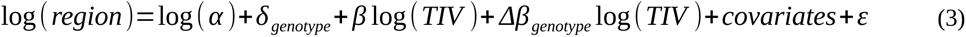

In equations (1–3), α and β represent the control scaling constant and scaling exponent, respectively, such that log(α) is the intercept and β the slope of the log-log relationship. The global reference was centered at the unrelated control mean, such that δ_genotype_ represents the regional difference at typical control brain size and was converted to a percentage as 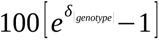.

In equation (3), Δβ_genotype_ represents deviation from the control scaling exponent. For SA and CT, TIV was replaced by total SA and mean CT, respectively. Because repeat sessions came only from carriers, the genotype-by-size interaction was estimated from between-participant differences in brain size (Supplementary Methods). Nested models were compared using likelihood-ratio tests on maximum-likelihood fits, and coefficients were estimated from restricted maximum-likelihood refits with Satterthwaite degrees of freedom ^(58, 59)^. To identify genotype-related allometric alterations, we conducted an omnibus screen comparing model (3) to model (1) separately for each CNV, using controls and carriers of that CNV. Regions with FDR-adjusted p < 0.05 were considered omnibus-positive. A positive screen could indicate a difference in the scaling constant or exponent. For volume, omnibus p-values were FDR-corrected separately within anatomical families (Supplementary Methods). For SA and CT, correction was applied across the 68 cortical parcels.

Within omnibus-positive regions, genotype-specific scaling exponents and constants were estimated from the joint model including all three groups, and a genotype-by-size interaction coefficient with nominal p < 0.05 indicated an altered scaling exponent, with Δβ quantifying the direction and magnitude of scaling-exponent deviation relative to controls.

As a post-hoc test to quantify overall scaling-exponent deviation, we calculated slope burden irrespective of omnibus significance. For each CNV, slope burden was defined as the mean absolute deviation from the control scaling exponent (mean |Δβ|), summarized descriptively across 98 non-redundant regional volume measures (68 cortical parcels, 29 aseg structures, and total corpus callosum volume) and tested formally across the 68 cortical regions for volume, SA, and CT. The test statistic was the deletion-minus-duplication difference in slope burden. Uncertainty was then estimated with a cluster bootstrap that resampled whole families with replacement and refit every regional model (2,000 draws), yielding the 95% percentile confidence interval and the proportion of draws in which deletion burden exceeded duplication burden.

### Integrating disproportionate and allometric phenotypes

Because the disproportion analysis estimates a single genotype effect under a common scaling exponent, it can be interpreted as an approximately fixed anatomical shift only where genotype-specific exponent deviation is small. We therefore masked the cortical disproportion map in regions with altered scaling exponents, defined first by a significant genotype-by-size interaction (p < 0.05) and, because this test has limited power, by deviation magnitude irrespective of significance (|Δβ| ≥ 0.3 and ≥ 0.2, derived from 10% and 5% tolerances for drift in regional size across the 10th–90th percentile TIV range and rounded to one decimal). To assess whether apparent exponent deviations exceeded estimation noise, Δβ estimates were also shrunk towards zero in proportion to their standard errors (empirical Bayes; Supplementary Methods; Fig. S8C).

## Results

### Sample characteristics

Demographic, imaging, clinical, and cognitive characteristics of the cohort are in Tables 1-2. Head circumference was above the population mean in deletion carriers and below it in duplication carriers. Mean full-scale IQ was lower in both CNV groups than in controls but did not differ between carrier groups. Autism was present in 18% of deletion and 10% of duplication carriers.

### Brain-size adjustment reveals disproportionate regional volume effects

Without TIV adjustment, covariate-adjusted models showed the expected dosage-dependent phenotype (Fig. 1A–C): relative to controls, deletion carriers had larger and duplication carriers smaller global (cGMV, sGMV, WMV, CC, SA) and regional volumes (median cortical d = +0.48 and −0.54; subcortical d = +0.34 and −0.85). Ventricular volume was the exception, increased in duplication (d = +0.32, FDR p = 0.024) but not deletion (d = +0.13, FDR p = 0.34).

**Figure 1:**
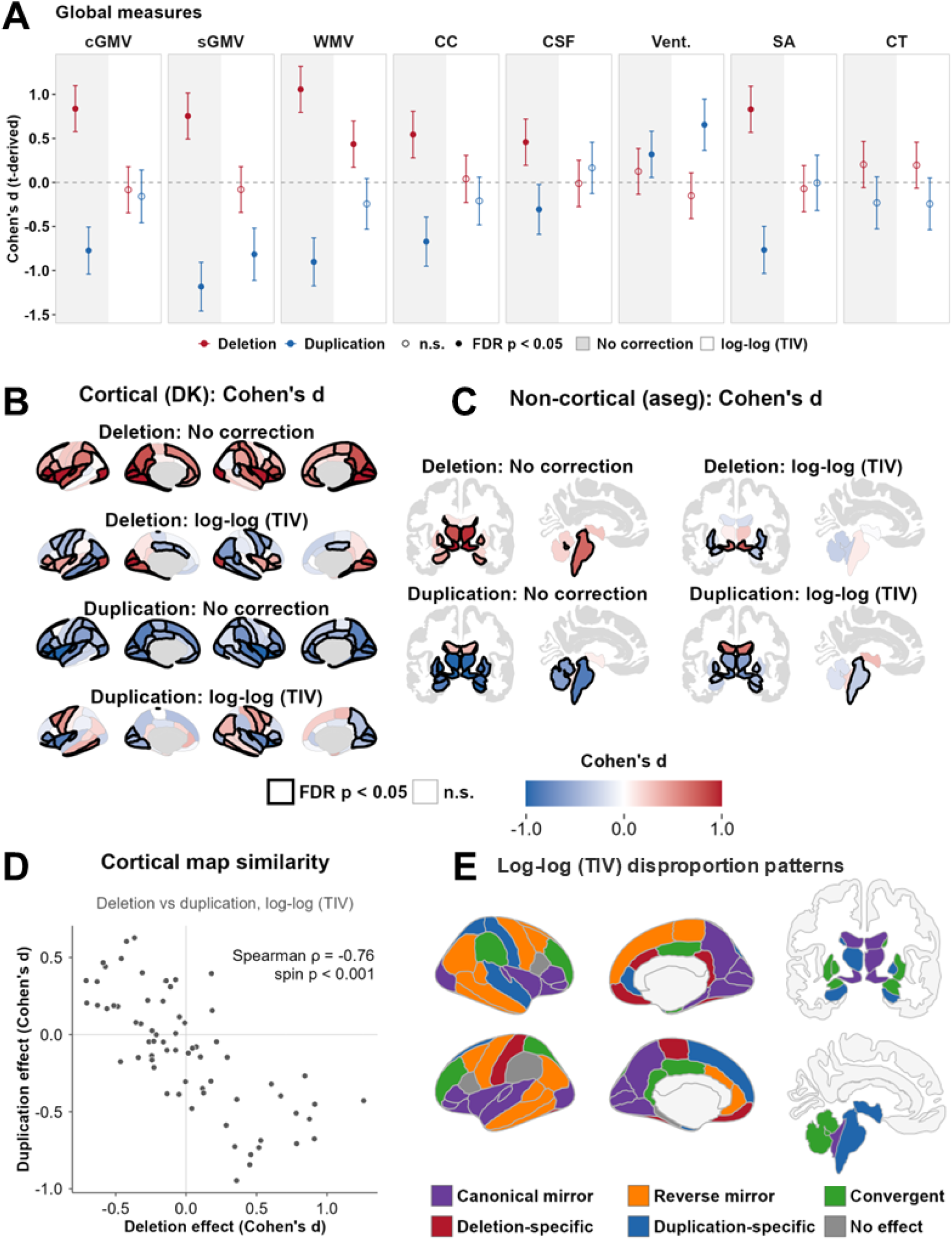
Brain-size adjustment reveals disproportionate regional volume effects in 16p11.2 copy-number variation. (A) Global measures shown as Cohen’s d for deletion and duplication relative to controls, before (shaded left half) and after (right half) log-log TIV adjustment. Filled points denote FDR p < 0.05 and open points nonsignificant effects; bars show approximate 95% confidence intervals. cGMV, cortical grey matter volume; sGMV, subcortical grey matter volume; WMV, white matter volume; CC, corpus callosum; CSF, cerebrospinal fluid; Vent., ventricles; SA, total surface area; CT, mean cortical thickness. (B) Cortical regional-volume effects in the DK atlas before and after log-log TIV adjustment. Red indicates larger and blue smaller volumes relative to controls; black outlines identify regions surviving FDR correction. (C) aseg-derived regional-volume effects, displayed as in (B). (D) Correspondence between log-log TIV-adjusted cortical deletion and duplication effects, one point per region; Spearman’s ρ with spherical spin-test significance. (E) Cortical and subcortical dosage-response patterns: canonical mirror (deletion enlargement, duplication reduction), reverse mirror (deletion reduction, duplication enlargement), convergent (same direction in both), or genotype-specific (effect in one genotype only). Effects with |d| ≤ 0.10 were treated as no effect (4 cortical regions).

Following log-log allometric adjustment for TIV, global volume differences were substantially attenuated (Fig. 1A). Only sGMV reduction (d = −0.82, FDR p = 2×10⁻⁶) and ventricular enlargement (d = +0.66, FDR p = 7×10⁻⁵) were disproportionate in duplication, and only WMV enlargement (d = +0.44, FDR p = 0.011) in deletion.

Regionally, adjustment had its largest impact in deletion (Fig. 1B, C), which showed bidirectional cortical disproportion, with 17 regions significantly larger and 17 smaller than expected for brain size. Duplication showed predominantly reductions (18 regions smaller, 7 larger), and all seven disproportionately enlarged regions were disproportionately reduced in deletion. Subcortical structures showed a different pattern. Deletion showed reductions in 5 of 21 structures (bilateral putamen and amygdala, left cerebellar WMV) and enlargement of the bilateral ventral diencephalon, whereas duplication showed reductions in 13 structures and no enlargement.

Deletion and duplication effect maps remained negatively correlated after adjustment, in both the cortex (before ρ = −0.75, after ρ = −0.76; both pspin < .001; Fig. 1D; Fig. S5A) and subcortex (ρ = −0.76 and −0.66; Fig. S5B, C). Dosage-response patterns, however, changed substantially (Fig. 1E; Fig. S5D, E). Before adjustment, cortical and subcortical effects were predominantly canonical mirror. After adjustment, the cortex remained predominantly reciprocal (40 of 68 regions: 22 canonical and 18 reverse mirror), whereas subcortical structures were dominated by convergent reductions (12 of 21; three canonical mirror, six duplication-specific).

Because previous neuroimaging studies of 16p11.2 CNVs accounted for brain size using proportional normalization or linear regression, we compared these approaches with our log–log model (Fig. S3). Linear regression produced regional volume effects highly similar to log–log adjustment (r > 0.9 in both genotypes; Fig. S3C, D), with subtle differences concentrated predominantly in hypoallometric occipital and sensory cortices (Fig. S3E, F). Proportional normalization recapitulated the overall pattern but deviated more from log–log estimates, particularly for subcortical structures in duplication (Fig. S3A–C). While regression-based approaches converged in this sample, only allometric adjustment estimates the scaling relationship as a parameter, allowing us to test whether 16p CNVs alter it.

### Distinct disproportionate phenotypes emerge in cortical surface area and thickness

We next asked whether the disproportionate cortical volume phenotype reflected alterations in cortical SA, CT, or both (Fig. 2). Before allometric adjustment, SA exhibited the expected reciprocal dosage pattern, with widespread enlargement in deletion (64 of 68 regions, 45 significant) and reductions in duplication (67 of 68 regions, 48 significant; Fig. 2A). CT differed: deletion showed few significant differences (six thicker, one thinner), whereas duplication showed predominantly thinning (19 thinner, 3 thicker; Fig. 2B).

**Figure 2.**
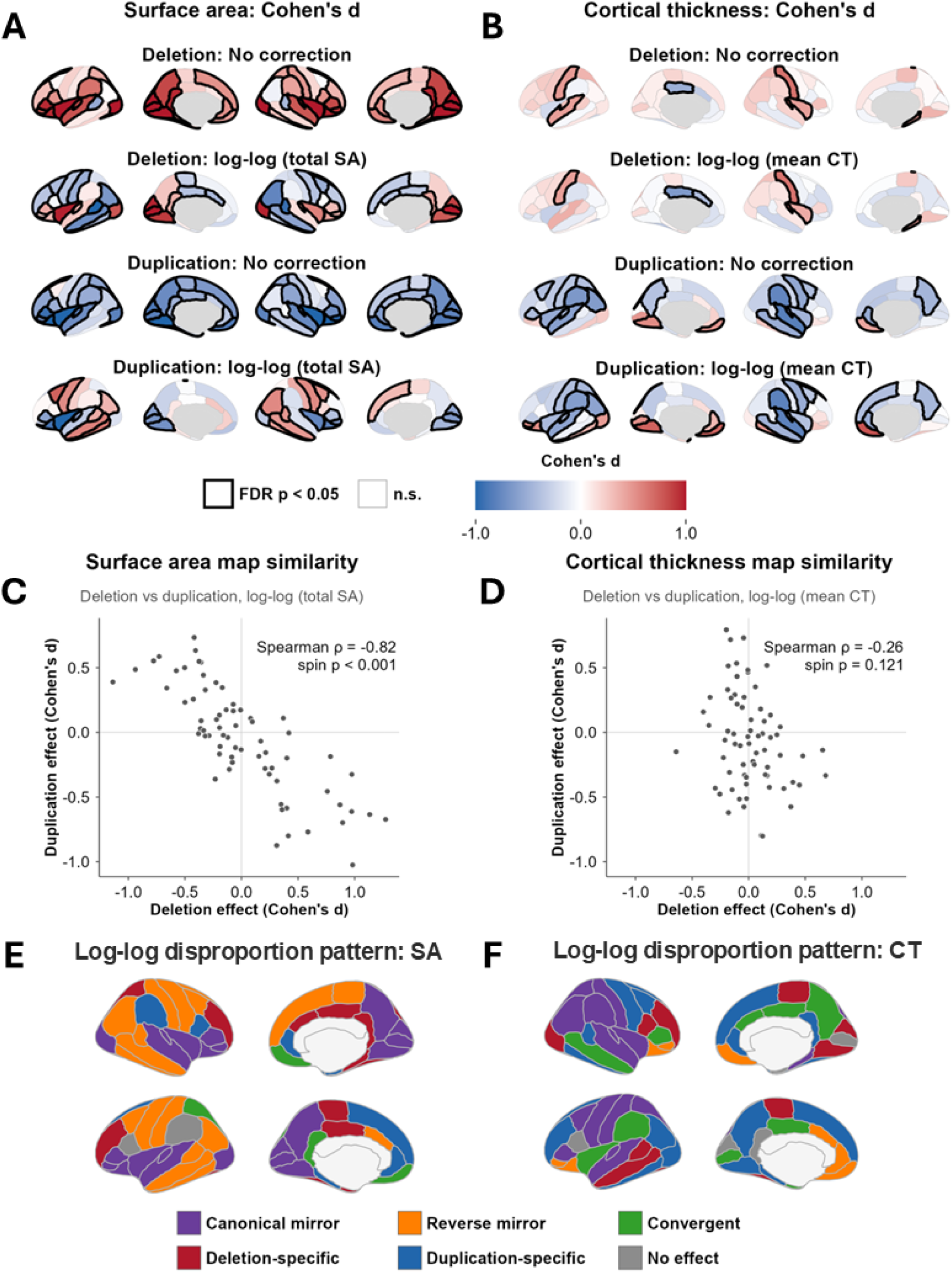
Distinct disproportionate cortical surface-area and cortical-thickness phenotypes in 16p11.2 copy-number variation. (A) Regional cortical SA effects before and after log-log adjustment for total cortical SA. Red indicates larger and blue indicates smaller regional SA relative to controls; black outlines identify regions surviving FDR correction (FDR p < 0.05). (B) Regional CT effects before and after log-log adjustment for mean CT, displayed as in (A). (C, D) Spatial correspondence between deletion and duplication effects on regional SA (C) and CT (D) after log-log adjustment; axes share the same ranges. (E, F) Adjusted SA (E) and CT (F) effects classified as in Fig. 1E.

Following log–log adjustment for total cortical SA, the regional SA phenotype closely paralleled the cortical disproportionate volume phenotype (Fig. 2A). Despite greater total SA, deletion carriers had widespread, bidirectional disproportion, with 19 regions significantly larger and 21 significantly smaller than expected. In duplication, adjustment revealed a more heterogeneous regional phenotype, with 16 regions significantly smaller and 13 significantly larger, despite no significant enlargements before adjustment. In contrast, adjustment for mean CT changed the CT phenotype little (deletion: five thicker, two thinner; duplication: 17 thinner, 11 thicker; Fig. 2B), with much smaller changes in regional effect size than for SA (mean |Δd| = 0.10 vs 0.52), indicating that regional CT differences were largely independent of mean CT.

Deletion and duplication SA effects were strongly negatively correlated both before and after adjustment (both ρ = −0.82, pspin < .001; Fig. 2C; Fig. S6A), whereas CT effects showed no significant correspondence (both ρ = −0.26, pspin = .12; Fig. 2D; Fig. S6B). Before adjustment, SA effects were predominantly canonical mirror (Fig. S6C) and CT effects predominantly duplication-specific (Fig. S6D). After adjustment, SA remained predominantly reciprocal but with substantially more reverse mirror effects (41 of 68 regions: 22 canonical, 19 reverse mirror; Fig. 2E), whereas CT remained heterogeneous (Fig. 2F), with 17 duplication-specific, 26 reciprocal (15 canonical, 11 reverse mirror), 10 convergent, 10 deletion-specific, and five negligible effects.

### Genotype-specific alterations in regional brain-size scaling

Having identified widespread regional disproportion, we next asked whether 16p CNVs also altered brain-size scaling itself (Fig. 3A). To do so, we conducted an omnibus screen for each CNV, comparing a common allometric model with one allowing genotype-specific scaling constants and exponents (eqs. 1 and 3). Omnibus effects were more widespread in deletion (51 of 105 volume measures, including 37 of 68 cortical regions) than in duplication (37 of 105, including 26 cortical regions; Fig. 3B). Among global measures, the screen was positive only for those showing significant disproportion (sGMV and ventricular volume in duplication, WMV in deletion; Fig. 1A), none of which showed a scaling-exponent difference (all p > 0.2).

**Figure 3:**
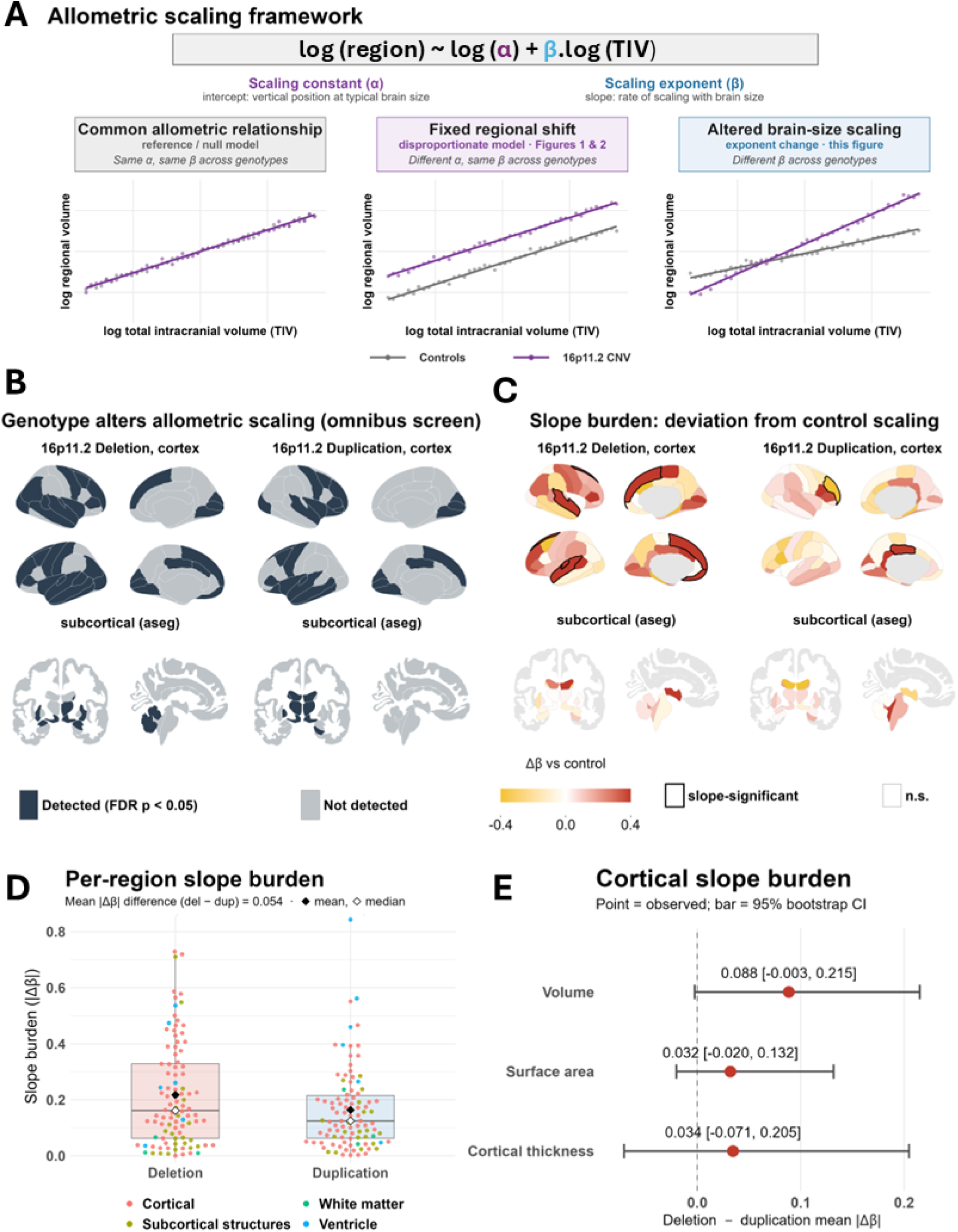
Genotype-specific alterations in regional brain-size scaling in 16p11.2 copy-number variation. (A) Schematic of a common allometric relationship, a fixed regional shift, and altered brain-size scaling; points are illustrative, not observed data. (B) Omnibus test for genotype differences in the scaling relationship between regional volume and TIV (scaling constant, slope, or both), tested separately for each CNV versus controls; dark regions survive FDR correction (FDR p < 0.05). (C) Genotype-specific difference in scaling slope relative to controls (Δβ); positive values indicate steeper and negative values shallower scaling. Black outlines identify omnibus-positive regions with a nominally significant slope difference (p < 0.05). Bilateral nucleus accumbens, slope-significant in deletion, is not shown in the aseg display atlas. (D) Absolute slope deviations (|Δβ|) across 98 non-redundant volume measures, colored by measure type, with box plots; white and black diamonds indicate medians and means. The deletion-duplication difference in mean |Δβ| is descriptive. (E) Deletion-duplication difference in mean |Δβ| across the 68 cortical regions for volume, SA, and CT; points show observed differences and bars show 95% intervals from 2,000 whole-family cluster bootstrap draws. Values above 0 indicate greater slope burden in deletion.

When we tested the genotype-by-size interaction directly (Fig. 3C), significant exponent differences were uncommon, occurring in 10 omnibus-positive measures in deletion (five prefrontal, three temporal, and the bilateral nucleus accumbens) and two in duplication. Among omnibus-positive measures without significant exponent differences, scaling-constant differences were detected in 37 of 41 deletion measures and all 35 duplication measures. Because the spatial pattern of exponent deviations suggested a broader deletion-dominant effect than per-region testing captured (Fig. 3C), as a post-hoc analysis, we also quantified the overall slope burden (mean |Δβ|) across measures.

Deletion showed a greater slope burden than duplication. Across 98 non-redundant regional volume measures, absolute slope deviations were greater in deletion for 54 measures with 0.054 higher mean slope burden (33% greater; Fig. 3D). The difference was largest in the cortex, where deletion showed greater deviations in 43 of 68 regions and a mean slope burden 0.088 higher than duplication (56% greater; 95% interval −0.003 to 0.215; 97.1% of draws favoring deletion; Fig. 3E). This suggests greater cortical scaling alteration in deletion, though simulations indicated limited power to detect a difference this large (60%; Supplementary Methods).

We next asked whether this pattern extended to SA and CT, testing each metric against its own global reference (regional maps in Fig. S7; burden differences in Fig. 3E). For SA, deletion showed more omnibus-positive regions (40 vs 27) and slope-significant regions (6 vs 1), greater absolute slope deviations in 41 of 68 regions, and a mean slope burden 0.032 higher than duplication (24% greater; 95% interval −0.020 to 0.132; 91.6% of draws favoring deletion). CT showed a different pattern: omnibus effects (25 vs 6) and slope-significant regions (7 vs 2) were more common in duplication, consistent with its largely duplication-specific CT phenotype. Mean CT slope burden was 0.034 higher in deletion (17% greater), but this difference was the least consistent across bootstrap draws (95% interval −0.071 to 0.205; 81.8% of draws favoring deletion).

### Integrating disproportionate and allometric phenotypes

Allometric TIV adjustment identified regional differences disproportionate to brain size (Figs. 1, 2), but interpreting these as fixed anatomical shifts assumes a shared scaling exponent. When exponents differ between genotypes, regional disproportion varies with brain size, so whether a disproportionate effect reflects a fixed shift depends on whether that region’s scaling exponent is altered.

Therefore, we masked the cortical disproportion map (Fig. 1) in regions with altered scaling exponents, defined first by significant exponent differences (Fig. 3C). However, the interaction test had limited power to detect exponent differences at a 10% tolerance (|Δβ| = 0.3; power 30% in deletion, 42% in duplication; Supplementary Methods). Thus, we next masked by magnitude, irrespective of significance (Fig. S8A, B): 24 deletion and 9 duplication cortical regions at the 10% tolerance (|Δβ| ≥ 0.3), and 35 and 19 at the 5% tolerance (|Δβ| ≥ 0.2). Empirical Bayes shrinkage revealed that duplication slope variation largely reflected noise, collapsing towards zero, whereas deletion retained a broader distribution (Fig. S8C).

Consequently, substantial exponent deviation affected fewer significantly disproportionate cortical regions in duplication (3 of 25, 12%) than in deletion (11 of 34, 32%). Importantly, altered scaling also occurred outside regions of significant disproportion: 7 of 8 deletion regions with significant exponent differences and both duplication regions were not significantly disproportionate (Figs. 1B, 3C). All significantly disproportionate subcortical structures remained unmasked under every rule (Fig. S8D). Regional allometric fits illustrate these distinctions for each measure (Fig. S9). Together, these findings suggest that disproportion in duplication predominantly reflects fixed shifts, whereas deletion more frequently shows altered scaling, both within and beyond regions of disproportion.

## Discussion

In this study, we examined how reciprocal 16p11.2 CNVs shape regional brain anatomy in the context of their opposing effects on overall brain size. Absolute neuroanatomical differences followed the well-established mirror pattern, but allometric TIV adjustment revealed widespread regional disproportion in both CNVs, driven primarily by surface area rather than cortical thickness. The spatial organization of this disproportion departed from the absolute mirror phenotype: cortical effects remained predominantly reciprocal, but nearly half of reciprocal regions showed the reverse mirror pattern, whereas subcortical structures showed predominantly convergent reductions in both CNVs. Allowing scaling exponents to vary further distinguished the two CNVs: regional disproportion in duplication largely reflected fixed shifts with preserved scaling exponents, whereas deletion showed more extensive scaling-exponent deviation, including in regions without significant disproportion. Thus, duplication largely preserved, whereas deletion more extensively disrupted, the scaling of regional anatomy with brain size

### Systematic regional analysis reveals diverse 16p11.2 dosage effects

Prior structural neuroimaging studies established the large reciprocal effects of 16p11.2 dosage on global brain anatomy alongside more heterogeneous regional effects ^(8–10)^. The earliest study described widespread increases in deletion and decreases in duplication spanning global and subcortical measures, but after intracranial volume normalization only the thalamus remained significant in both CNVs, leading the authors to conclude that 16p11.2 exerts a pervasive rather than selective effect on the brain ^(8)^. Yet, subsequent studies identified additional effects beyond brain size, including grey matter volumes that varied inversely with copy number across cortical and subcortical regions ^(9)^ and regions preferentially affected by one CNV, such as the calcarine and transverse temporal cortices in deletion and the caudate and hippocampus in duplication ^(10)^.

Using the Simons VIP cohort, which was also the sample used in most prior studies ^(8–10)^, we systematically characterized the joint regional effects of deletion and duplication while explicitly modeling the expected allometric relationship between brain size and regional volume. Allometric TIV adjustment revealed widespread regional disproportion in both CNVs. Cortical disproportion remained predominantly reciprocal but included nearly equal canonical and reverse mirror patterns with distinct spatial distributions: canonical mirror effects clustered in primary visual and auditory, insular, and ventrolateral prefrontal cortex, whereas reverse mirror effects clustered in dorsal frontal, sensorimotor, and lateral temporal and parietal association cortex. Deletion-specific effects were predominantly reductions in ventral temporal and medial frontal cortex, whereas duplication-specific effects were sparse and spatially scattered. In the subcortex, convergent reductions involved the basal ganglia, amygdala, and cerebellum, whereas duplication-specific reductions involved the hippocampus and thalamus.

The more extensive regional phenotype we identified could reflect differences in analytic approach, particularly how brain size and age were modeled. For example, Qureshi et al. (2014) used proportional normalization and found few effects beyond brain size ^(8)^. In contrast, Martin-Brevet et al. (2018) additionally examined the cortex at the voxel and vertex level using regression-based linear adjustment, identifying more extensive regional effects beyond those expected for brain size, consistent with the phenotype observed here ^(10)^. This parallels our sensitivity analyses, in which proportional normalization diverged substantially from regression-based estimates, particularly in the subcortex, whereas linear and log-log adjustment produced highly similar results ^(17–19, 30, 67)^. This concordance does not imply that regional anatomy scales linearly with brain size; rather, linear adjustment approximates the allometric relationship in this sample. Although, these approaches may diverge across broader ranges of brain sizes. Differences between studies may also reflect age modeling, anatomical parcellation, image processing and quality control, scanner harmonization, and treatment of familial and repeated observations ^(8, 10)^.

### Surface area and cortical thickness reveal distinct components of the cortical phenotype

Examining SA and CT further distinguished the effects of 16p11.2 dosage. The disproportionate cortical volume phenotype was driven predominantly by SA, consistent with earlier evidence that cortical volume differences in 16p CNVs primarily reflect SA rather than thickness ^(8)^. Regionally, SA effects closely recapitulated cortical volume, including both canonical and reverse mirror patterns. CT showed a markedly different phenotype, with less extensive disproportionate effects that were concentrated predominantly in duplication. Prior studies have similarly reported non-mirror organization of CT, with thickness reduced in both deletion and duplication while global volume and SA varied with copy number ^(9)^. Thus, despite their reciprocal effects on overall brain size and cortical SA, deletion and duplication do not exert simply opposing effects across all components of cortical morphology.

The predominance of surface-area effects may also point to developmental processes underlying the cortical phenotype. SA and CT have partly distinct developmental determinants ^(20, 46)^, with cortical surface expansion linked to progenitor proliferation ^(44, 68, 69)^. Patient-derived neural precursor cells carrying the 16p11.2 deletion show hyperproliferation and altered mitogenic responses, providing a potential cellular link to altered cortical surface expansion ^(40)^. Reciprocal-CNV models likewise show both opposing and shared cellular effects, including reciprocal differences in neuronal morphology and organoid growth but convergent reductions in synaptic density and neuronal migration ^(38, 39)^. Given the distinct developmental determinants of SA and CT, the divergence between the predominantly reciprocal SA phenotype and the duplication-weighted thickness phenotype raises the possibility that deletion and duplication affect cortical anatomy through partly distinct developmental mechanisms ^(45, 46, 70)^. However, mechanistic studies remain weighted towards deletion, leaving the cellular basis of the duplication phenotype comparatively unexplored.

### 16p11.2 deletion more extensively alters conserved allometric scaling

Beyond identifying regional disproportion, the allometric framework allowed us to test whether 16p11.2 dosage alters scaling itself. Regional brain volumes follow characteristic scaling relationships with overall brain size that reflect conserved developmental and evolutionary constraints on brain architecture ^(13, 16, 33–35, 37, 71, 72)^. These relationships are highly reproducible across populations and largely stable across age and sex ^(15, 31, 32)^. This conservation holds despite substantial individual variation in regional size: individuals differ in regional volume while sharing the same scaling exponent. Genetic variants that alter brain size could therefore act in two ways: by shifting regional size within conserved scaling, as in normal variation, or by changing the scaling exponent itself, a more fundamental departure in which regional differences depend on brain size.

This distinction is particularly important in 16p CNVs, where large reciprocal differences in brain size could otherwise obscure which form genotype-associated regional differences take.

Notably, deletion and duplication differed in whether their regional differences reflected fixed shifts or altered scaling exponents. In duplication, regional disproportion largely reflected fixed shifts: substantial exponent deviation affected only 12% of disproportionate cortical regions, and after empirical Bayes shrinkage, apparent duplication slope deviations collapsed towards zero, consistent with estimation noise. In deletion, exponent deviation was more extensive, affecting 32% of disproportionate cortical regions and persisting after shrinkage. It also extended beyond regions of disproportion: most significant deletion exponent differences, including in bilateral superior frontal, frontal pole, and superior temporal cortex, occurred in regions that were not disproportionate at typical brain size. Although individually significant exponent differences were uncommon, reflecting the limited power of region-level tests, a post-hoc analysis showed greater aggregate cortical slope burden in deletion (97.1% of bootstrap draws), more consistently for SA (91.6%) than CT (81.8%), suggestive of more extensive scaling alteration in deletion. Thus, duplication disproportion largely preserved scaling exponents, whereas deletion altered scaling both within and beyond regions of disproportion.

The more extensive scaling-exponent deviation in deletion is especially notable because duplication, despite its marked effect on overall brain size in the opposite direction, largely preserved normative scaling exponents. Prior studies also suggest that such alterations are not an inevitable consequence of altered brain size or neurodevelopmental disorder. In sex chromosome aneuploidies, regional volumes largely conformed to normative allometric predictions, with deviations limited to select structures, consistent with largely preserved scaling, although scaling exponents were not tested directly ^(28, 29)^. Similarly, a large study of idiopathic autism that directly compared scaling relationships found no robust differences in subcortical scaling ^(73)^. Thus, the more extensive scaling-exponent deviation in 16p11.2 deletion may represent a distinctive feature of its neuroanatomical phenotype. Alternatively, because scaling exponents have rarely been tested directly, altered scaling may be an underexplored phenotype across neurodevelopmental conditions that alter brain size.

The dissociation in scaling patterns between deletion and duplication is particularly striking given the reciprocal gene dosage and strongly mirrored effects on overall brain size and absolute regional anatomy. Together with the divergent SA and CT phenotypes, it suggests that deletion and duplication influence regional brain organization through partly distinct developmental mechanisms rather than perturbing the same processes in opposite directions. This is consistent with cellular studies that have identified both opposing and shared effects across developmental phenotypes in deletion and duplication ^(38, 39)^. Thus, the divergent allometric effects observed here may provide a macroscale signature of similarly non-reciprocal developmental processes.

## Limitations

Despite the relatively large imaging sample for a rare CNV, carrier numbers remain modest, particularly for estimating genotype-by-brain-size interactions. Individually significant scaling-exponent differences were therefore uncommon (estimated power of 30% in deletion and 42% in duplication to detect |Δβ| = 0.3), making the post-hoc burden analysis a useful complement for assessing systematic genotype differences across regions. Simulations indicated limited power (60%) to detect a cortical burden difference of the observed size, so the deletion–duplication burden difference should be regarded as suggestive. Nevertheless, formally testing this difference, rather than relying on visual comparison of exponent maps as is common, provides a calibrated estimate of its uncertainty. Replication in larger independent samples will be important, as Simons VIP may not capture the full spectrum of neuroanatomical variation associated with 16p CNVs. Age differed across genotype groups, with deletion carriers younger than duplication carriers and controls. Although age was modeled flexibly with splines, developmental differences in regional brain-size scaling could contribute to genotype-specific estimates. Despite harmonization, quality control, and adjustment for scanner effects, the multisite cohort may contribute residual acquisition-related heterogeneity.

## Conclusion

In sum, the neuroanatomical consequences of 16p11.2 dosage extend beyond reciprocal changes in overall brain size. Allometric adjustment revealed distinct patterns of regional disproportion, and direct examination of scaling distinguished the two CNVs: in duplication, regional disproportion predominantly reflected fixed shifts with preserved scaling exponents, whereas in deletion, scaling-exponent deviation was more extensive, both within and beyond regions of disproportion. Thus, reciprocal 16p11.2 dosage does not simply shift regional anatomy in opposite directions: duplication largely preserves the normally conserved scaling of regional anatomy with brain size, whereas deletion more extensively disrupts it, suggesting that reciprocal gene dosage may produce nonreciprocal effects on neurodevelopment.

## Data & Code Availability

### Data availability

Simons Searchlight (formerly Simons VIP) data are available to qualified researchers through SFARI Base (https://base.sfari.org). Under the SFARI data use agreement, the participant-level data used in these analyses cannot be shared by the authors. Summary statistics for all regional analyses are provided in the Supplementary Information.

### Code availability

Analysis code will be made publicly available upon publication at https://github.com/BGDlab/16p11.2-allometry.

