## Supplemental Methods for "Reciprocal 16p11.2 Copy Number Variants Differentially Alter Regional Brain Anatomy and Allometric Scaling"

#### Participants:

Participants were drawn from the Simons Variation in Individuals Project (Simons VIP), a genetics-first cohort that recruited 16p11.2 deletion and duplication carriers without regard to clinical diagnosis or age [(1)]. Eligibility required the canonical ~600-kb 16p11.2 deletion or duplication, or a smaller CNV at the locus, confirmed by fluorescent in situ hybridization or comparative genomic hybridization; cascade testing identified additional carriers within families. Individuals with other pathogenic CNVs or with neurogenetic or neurological diagnoses unrelated to 16p11.2 were excluded. Non-carrier family members were enrolled as familial controls. Unrelated controls were excluded if they had a major psychiatric diagnosis or a family member with a neurodevelopmental disorder or genetic abnormality [(2)]. Structural MRI was acquired only in participants able to complete scanning without sedation.

#### MRI Acquisition Parameters:

Research scans were acquired at University of California sites and the Children's Hospital of Philadelphia on matched 3-T Siemens Magnetom TIM Trio scanners. Research multi-echo MPRAGE scans were acquired using 176 slices, a 256 × 256 matrix, repetition time = 2530 ms, inversion time = 1200 ms, first echo time = 1.64 ms, and flip angle = 7°. Echoes were combined using root-mean-square averaging. Clinical scans were acquired at the University of Washington Medical Center, Baylor University Medical Center, and Boston Children's Hospital on 3-T Philips Achieva or Siemens Magnetom TIM Trio scanners. Clinical single-echo MPRAGE scans were acquired using 160 slices, a 256 × 256 matrix, echo time = 2.98 ms, repetition time = 2300 ms, and flip angle = 9°.

#### Anatomical measures and region definitions

Global measures were obtained from the FreeSurfer global outputs. Cortical analyses included the 68 regions of the Desikan-Killiany (DK) atlas. Two anatomical sets were used for the non-cortical analyses. The subcortical set comprised 21 measures: bilateral hippocampus, amygdala, caudate, putamen, pallidum, thalamus, accumbens area, ventral diencephalon, cerebellar cortex, and cerebellar white matter, plus the brainstem. The aseg set comprised these 21 structures plus the lateral, inferior lateral, third, and fourth ventricles and hemispheric cerebral white matter (n = 29). Total corpus callosum volume was analyzed as a single composite rather than its five FreeSurfer subdivisions, whose internal boundaries partition a continuous structure, and was reported with global measures.

#### Analytic sample preparation

Only single- or multi-echo MPRAGE sessions with known diagnostic group and age and no missing morphometric measures were included. Non-positive regional volumes, which occasionally result from ComBat harmonization, were excluded from the corresponding regional model. Triplication carriers (n = 2; 4 sessions) were excluded. Two additional participants were excluded: one deletion carrier with a structural brain abnormality on MRI and one duplication carrier with an additional CNV outside 16p11.2. Corpus callosum measures were set to missing for one participant with a corpus callosum segmentation failure. When multiple eligible acquisitions were available for the same session, multi-echo MPRAGE was prioritized; within acquisition type, the scan with the highest SynthSeg+ QC score was retained. All eligible longitudinal sessions were included, with dependence among repeated observations addressed in the statistical analyses.

**Integrating disproportionate and allometric phenotypes**

To determine where scaling-exponent deviation limited interpretation of the disproportionate estimate, regions with altered scaling were masked from the disproportionate maps. We first masked regions with a detected scaling-exponent difference, defined as omnibus-positive regions with a nominal genotype-by-size interaction ($p < 0.05$). Because per-region interaction tests have limited power to detect the distributed exponent deviation assessed by the slope-burden analysis, we additionally applied an effect-size threshold to all regions, independent of statistical significance and omnibus status. To quantify this limitation, power of the interaction test was estimated for each cortical region from the standard error of its interaction coefficient, assuming a two-sided test at $\alpha = 0.05$ and a true exponent difference of $|\Delta\beta| = 0.3$, and averaged across the 68 cortical regions (30% in deletion, 42% in duplication). Thresholds were defined by the tolerated variation in the estimated genotype effect across the observed 10th-to-90th-percentile TIV range; for 5% variation:

$$|\Delta\beta| < \frac{\log(1.05)}{\log(TIV_{90}/TIV_{10})} = 0.166$$

The primary threshold allowed ≤10% variation ($|\Delta\beta| < 0.3$) and a stricter threshold allowed ≤5% variation ($|\Delta\beta| < 0.2$); thresholds were rounded to one decimal place from the exact values (0.325 and 0.166).

To assess whether apparent exponent deviations exceeded estimation noise, regional $\Delta\beta$ estimates for each CNV were subjected to empirical Bayes shrinkage [3-5]. True regional slope differences were modeled as draws from a normal distribution with mean zero and variance $\tau^2$, estimated separately for cortical regions and for all other measures by the method of moments as the mean of $\Delta\beta^2 - SE^2$ (truncated at zero), where SE denotes the standard error of each regional estimate. Each estimate was then shrunk toward zero by the factor $\tau^2/(\tau^2 + SE^2)$, such that less precise estimates were shrunk more strongly (Fig. S8C). Shrinkage was applied in closed form to the existing regional estimates, without refitting the regional models, and assumes independence among regional estimates. For subcortical structures, the 5% threshold was additionally applied to the shrunken estimates (Fig. S8D).

We additionally characterized the scaling constant among omnibus-positive regions without a nominally significant genotype-by-size interaction. In these regions, a genotype coefficient of $p < 0.05$ in the genotype-specific constant model indicated an altered scaling constant, representing a regional difference approximately fixed across the sampled brain-size range.

### Additional Statistical Details:

*Disproportionate Analyses:*

Mixed-effects models were fit by restricted maximum likelihood, with Satterthwaite degrees of freedom used for fixed-effect inference. FDR correction was performed separately by model and genotype contrast within four families: the eight whole-brain tissue compartments displayed in Figure 1A, 68 cortical parcels, 29 aseg measures, and remaining modeled global measures. Global measures were corrected separately so that composites could not influence significance among the regional measures they summarize. For surface area and cortical thickness analyses, FDR correction was performed across the 68 cortical parcels separately by metric, model, and genotype contrast.

To assess spatial correspondence between deletion and duplication volume, surface area, and cortical thickness effect-size maps before and after allometric adjustment, we calculated Spearman correlations. For cortical maps, significance was evaluated using 5,000 spherical rotations of Desikan–Killiany parcel centroids on the fsaverage sphere. Subcortical correspondence was summarized by Spearman's ρ without a significance test. Spin tests were two-sided; empirical lower- and upper-tail probabilities were calculated with the +1 correction, and the p value was defined as twice the smaller tail probability, capped at 1.

*Sensitivity analysis of brain-size adjustment methods:*
To assess sensitivity of regional effects to brain-size adjustment strategy, we compared unadjusted, proportional, linear, and log–log models across regional volume, surface area, and cortical thickness analyses. Agreement between adjustment approaches was quantified within genotype and anatomical class using Pearson correlations between regional Cohen's *d* estimates; 95% confidence intervals were calculated using Fisher's *r*-to-*z* transformation.

We further characterized differences between linear and log–log adjustment by calculating the regional effect-size difference, $\Delta d = d_{linear} - d_{\log-\log}$. Cortical volume and surface-area regions with $|\Delta d| > 0.10$ in either genotype were considered meaningfully divergent for descriptive follow-up. Within these regions, linear and log–log models were refit and population-level predictions were obtained from fixed effects. Log–log predictions were retransformed to the native measurement scale using Duan's smearing correction. For each observation and region, prediction divergence was calculated as the difference between log–log and linear predictions divided by the standard deviation of the regional measure. Subject-level divergence was summarized as the mean absolute standardized prediction difference across divergent regions.

*Global reference sensitivity analyses.*
To evaluate the choice of global reference measure for cortical surface area and thickness, we compared the variance explained by the phenotype-specific global measure with that explained by TIV. Among controls, partial $R^2$ was calculated for each of the 68 cortical parcels comparing total SA with TIV for regional SA and mean CT with TIV for regional CT, adjusting for age spline, sex, and mean bilateral Euler number. Sensitivity analyses then repeated the log-log regional SA and CT models using TIV rather than total SA or mean CT as the global reference (Fig. S4).

*Altered brain-size scaling analyses:*
The omnibus screen was conducted separately for each CNV: model (3), containing genotype-specific scaling constants and exponents, was compared with model (1), containing neither

genotype term, using a likelihood-ratio test on maximum-likelihood fits, in controls and deletion carriers and in controls and duplication carriers. For each CNV, a region was considered omnibus-positive if its CNV-specific screen reached FDR-adjusted $p < 0.05$, with FDR correction performed separately within four families: the 68 cortical parcels, the 29 aseg measures, the five global measures (cGMV, sGMV, WMV, ventricular volume, and total corpus callosum volume), and the remaining composite measures (total grey matter volume and left and right cortical volume).

Slope-burden uncertainty was estimated with a cluster bootstrap of 2,000 draws. In each draw, families were resampled with replacement from all families, with each family's participants and sessions kept intact, and every regional model was refit with the full mixed-effects specification; fits with a singular family variance component or a convergence warning were retained. For volume, SA, and CT separately, the deletion-minus-duplication difference in mean $|\Delta\beta|$ across the 68 cortical parcels was recalculated in each draw. We report the observed difference, its 95% percentile interval (2.5th and 97.5th percentiles of the bootstrap distribution), and the percentage of draws in which the difference was positive. Volume burden across a 98-measure nonredundant set, which excluded composite measures whose components were already represented, is reported descriptively; total corpus callosum volume was retained in this set because its component subdivisions were not modeled separately.

Power and false-positive rates of the cortical volume burden test were estimated by simulation. Simulated datasets retained the real sample's participants, families, sessions, brain size, age, sex, and Euler number; regional outcomes were generated from the fitted models, with family, participant, and session variance components and inter-regional residual correlations taken from the real data, and with deletion and duplication slope deviations set by scenario. For power, slope deviations were set to the observed deletion and duplication estimates (true burden difference 0.088; 100 datasets). In three null scenarios with equal burden (200 datasets each), deletion and duplication shared the same slope deviation in every region, or the deletion map, rescaled to the average observed burden, was assigned to duplication either in shuffled regions or with reversed sign. In each dataset, variance components were estimated once and held fixed while the burden difference was evaluated with 499 family-level bootstrap draws; a difference was considered detected when its 95% percentile interval excluded zero. Power was 60% (95% CI 50–69%), and false-positive rates ranged from 4.0% to 6.5%.

*Repeat-session sensitivity analyses*

In the primary allometric models, each participant's global reference was split into the participant's mean across sessions and each session's deviation from that mean. The genotype-by-size interaction was specified on the participant-mean component, so that altered scaling exponents were estimated from between-participant differences in brain size, because repeat sessions came only from carriers. Two sensitivity analyses instead used one observation per participant: the earliest session, or log-transformed regional and global measures, age, and Euler number averaged across each participant's sessions. Because each participant contributed a single observation, these models included a random intercept for family only.

Results were consistent with the primary analysis: regional cortical volume slope differences were highly correlated with primary estimates (r = 0.97–0.995), the same numbers of cortical regions showed significant exponent differences (8 in deletion, 2 in duplication), and cortical slope burden remained greater in deletion than duplication for volume, SA, and CT in both analyses.
