## Supplemental Figures for "Reciprocal 16p11.2 Copy Number Variants Differentially Alter Regional Brain Anatomy and Allometric Scaling"

### Figure S1: Image Quality Metrics by Site and Genotype

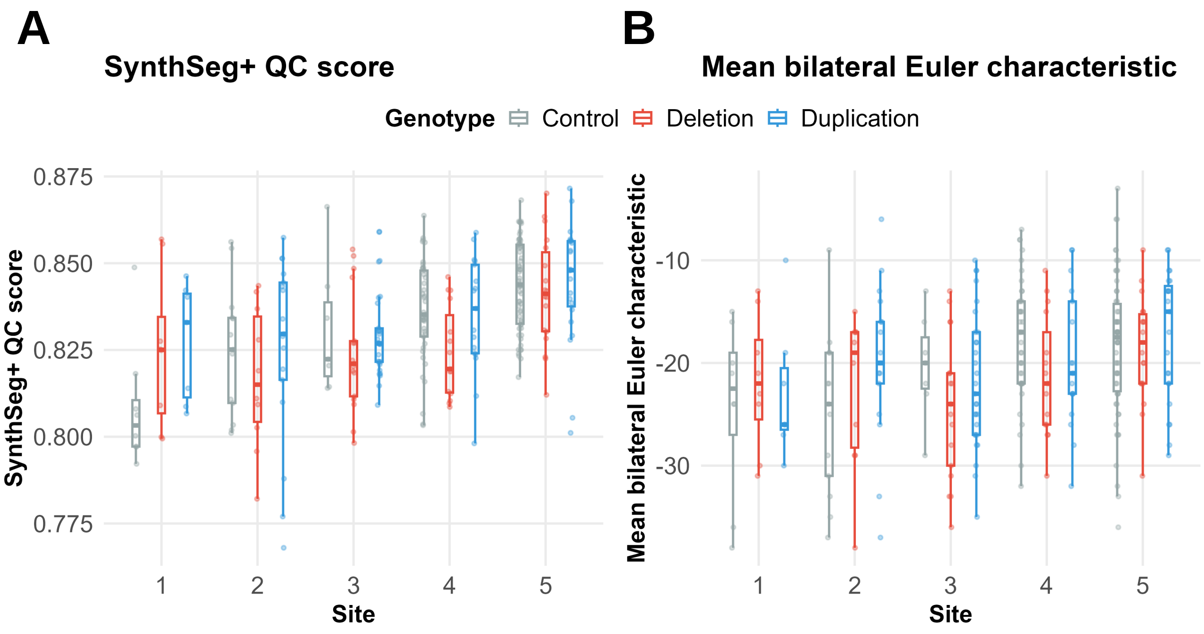


**Supplementary Figure S1: Image-quality metrics by site and genotype.** Distributions of (A) SynthSeg+ quality-control (QC) scores and (B) mean bilateral Euler characteristic in the final post-QC analytic sample, shown by acquisition site and genotype. Boxplots indicate the median and interquartile range, with whiskers extending to 1.5× the interquartile range; points represent individual imaging sessions. Familial and unrelated noncarrier controls are combined as controls. Sites are numbered 1-5 to maintain anonymity.

### Supplementary Figure S2. Raw volume distributions — cortical (1/5)

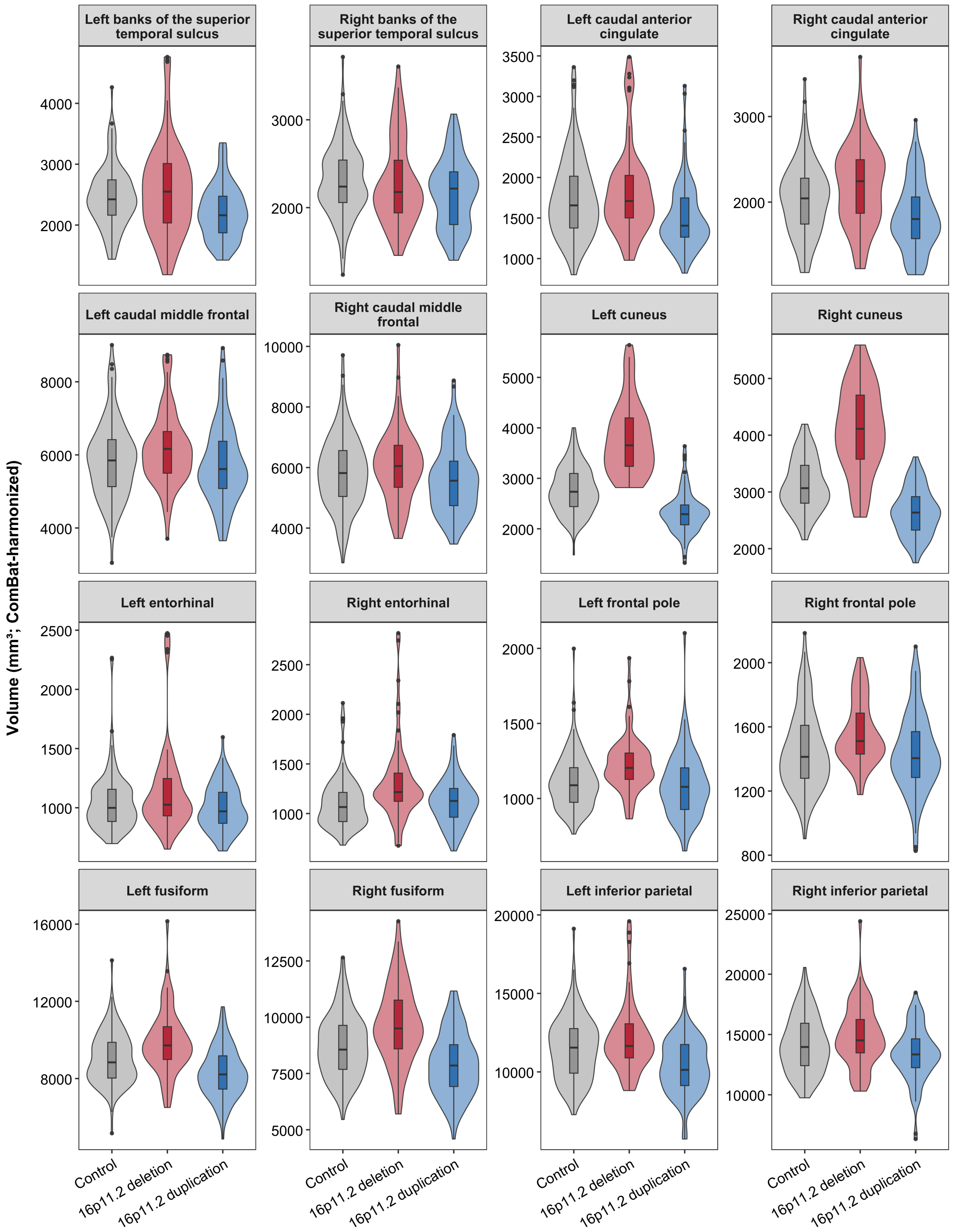

### Supplementary Figure S2. Raw volume distributions — cortical (2/5)

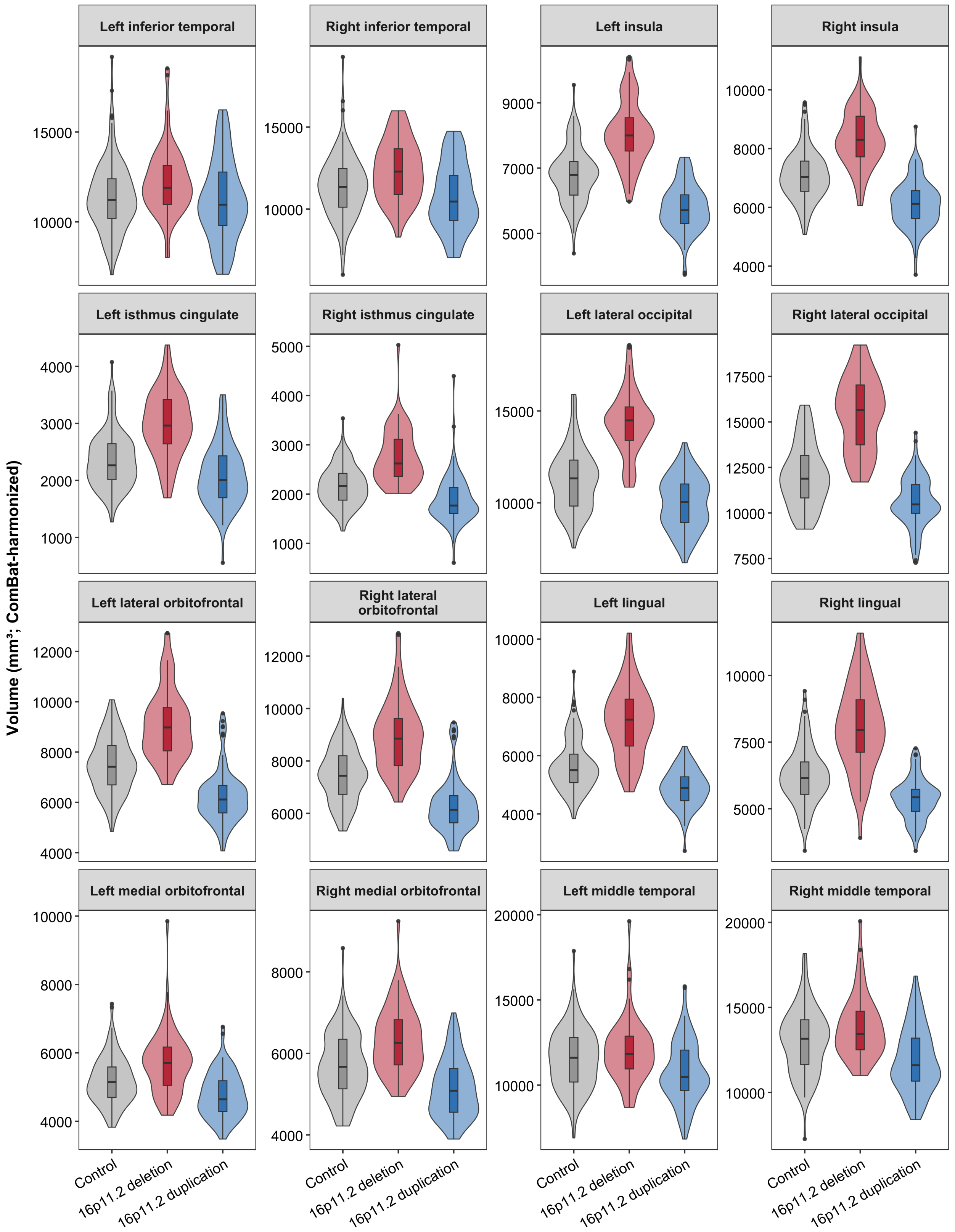

### Supplementary Figure S2. Raw volume distributions — cortical (3/5)

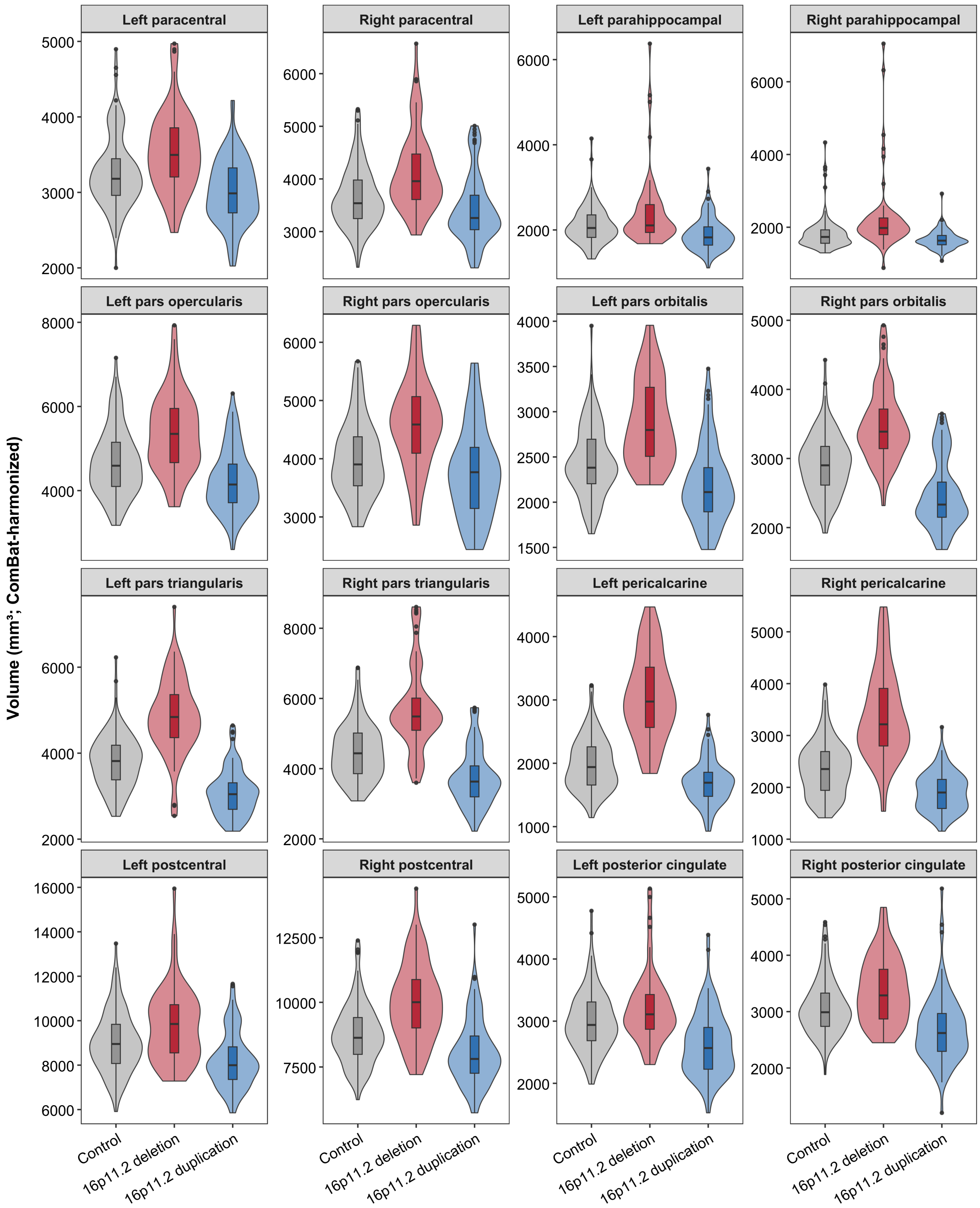

### Supplementary Figure S2. Raw volume distributions — cortical (4/5)

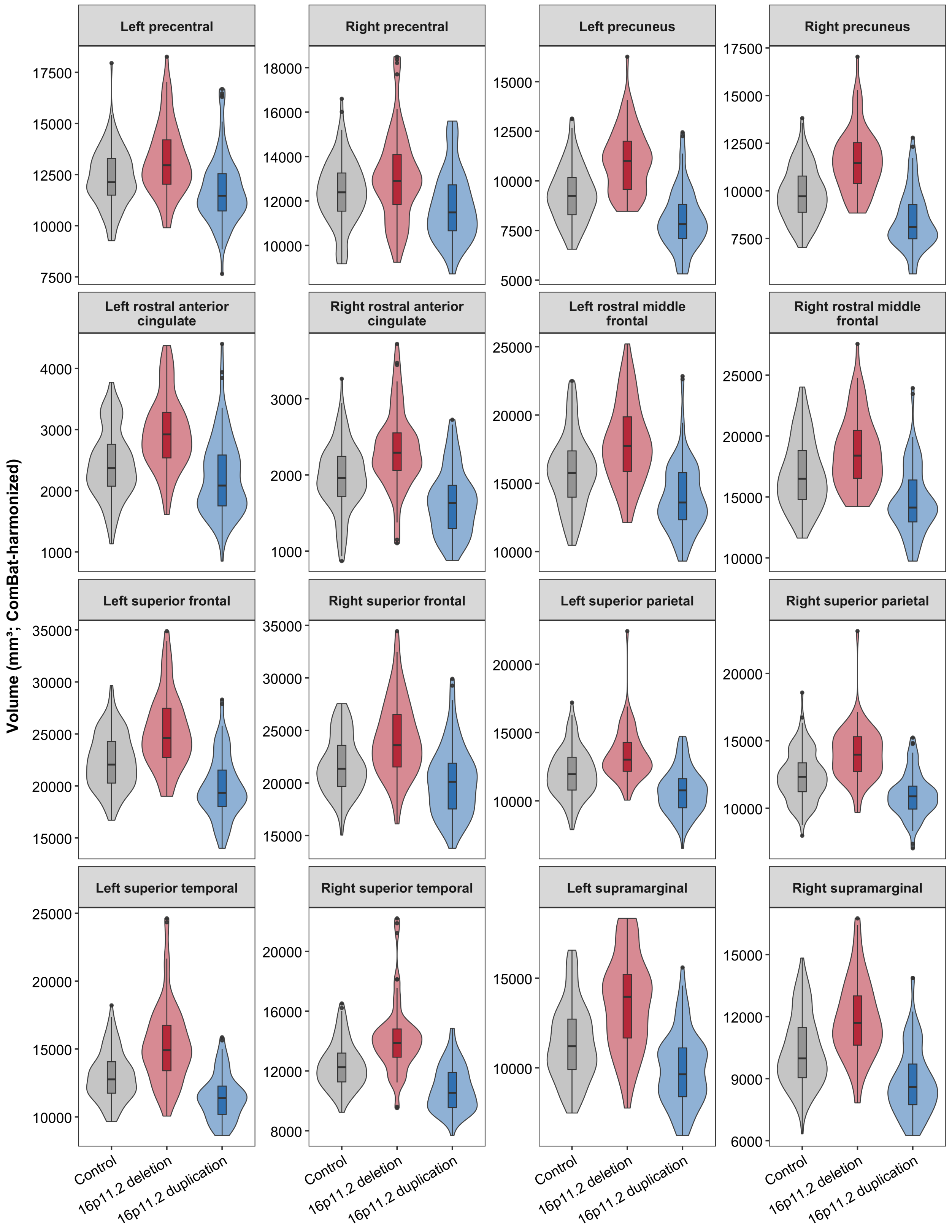

### Supplementary Figure S2. Raw volume distributions — cortical (5/5)

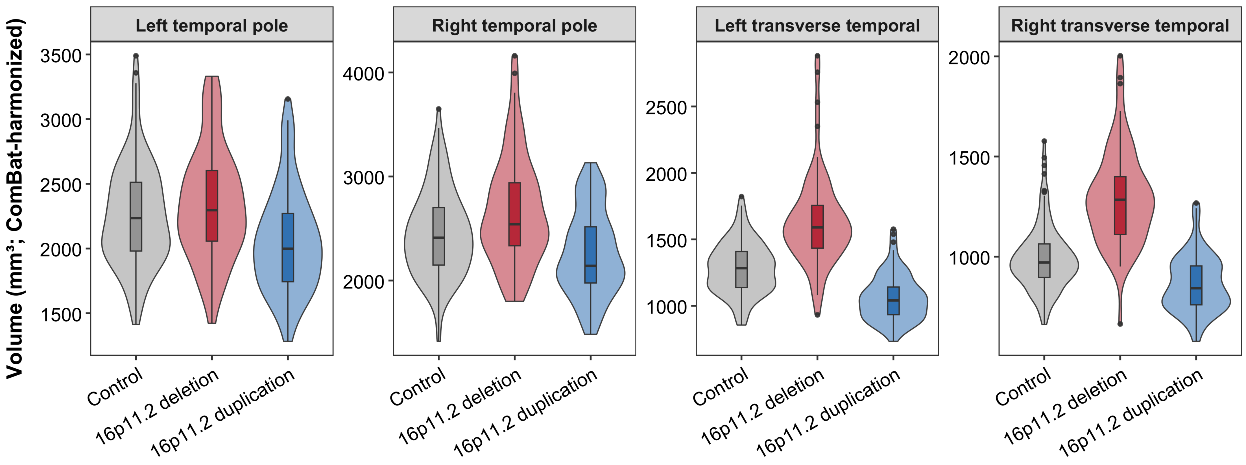

### Supplementary Figure S2. Raw volume distributions — subcortical

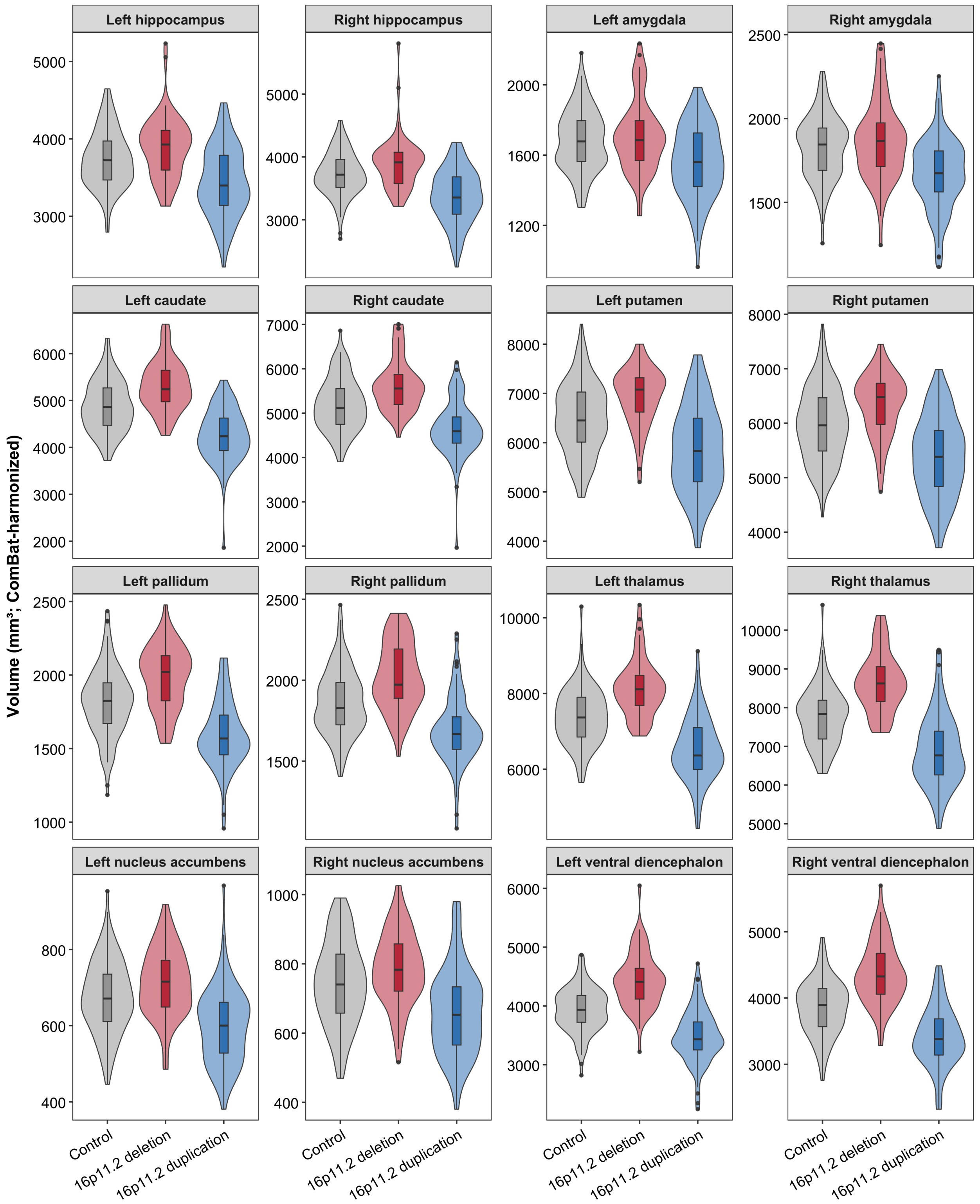

### Supplementary Figure S2. Raw volume distributions — global & other

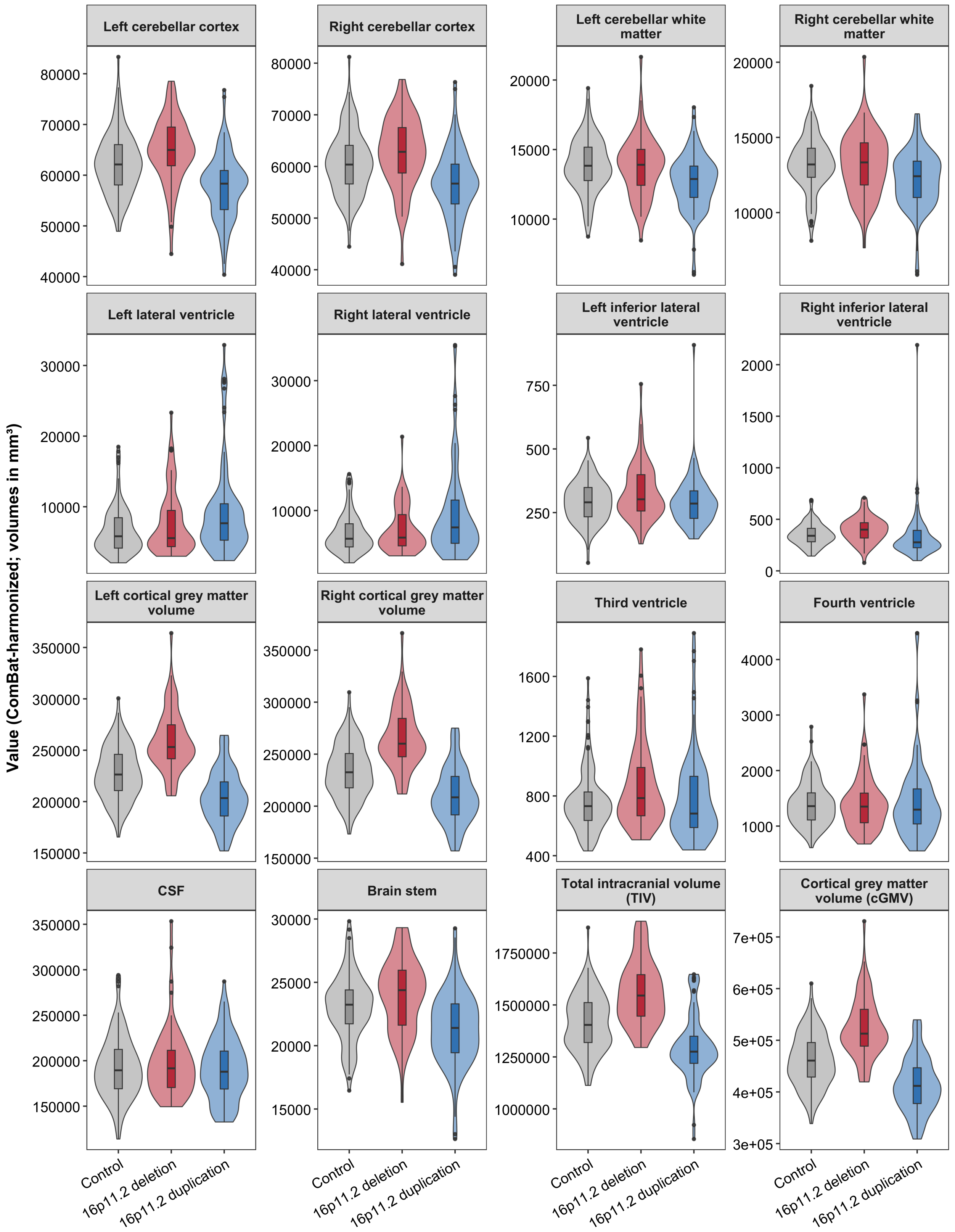

### Supplementary Figure S2. Raw volume distributions — global & other

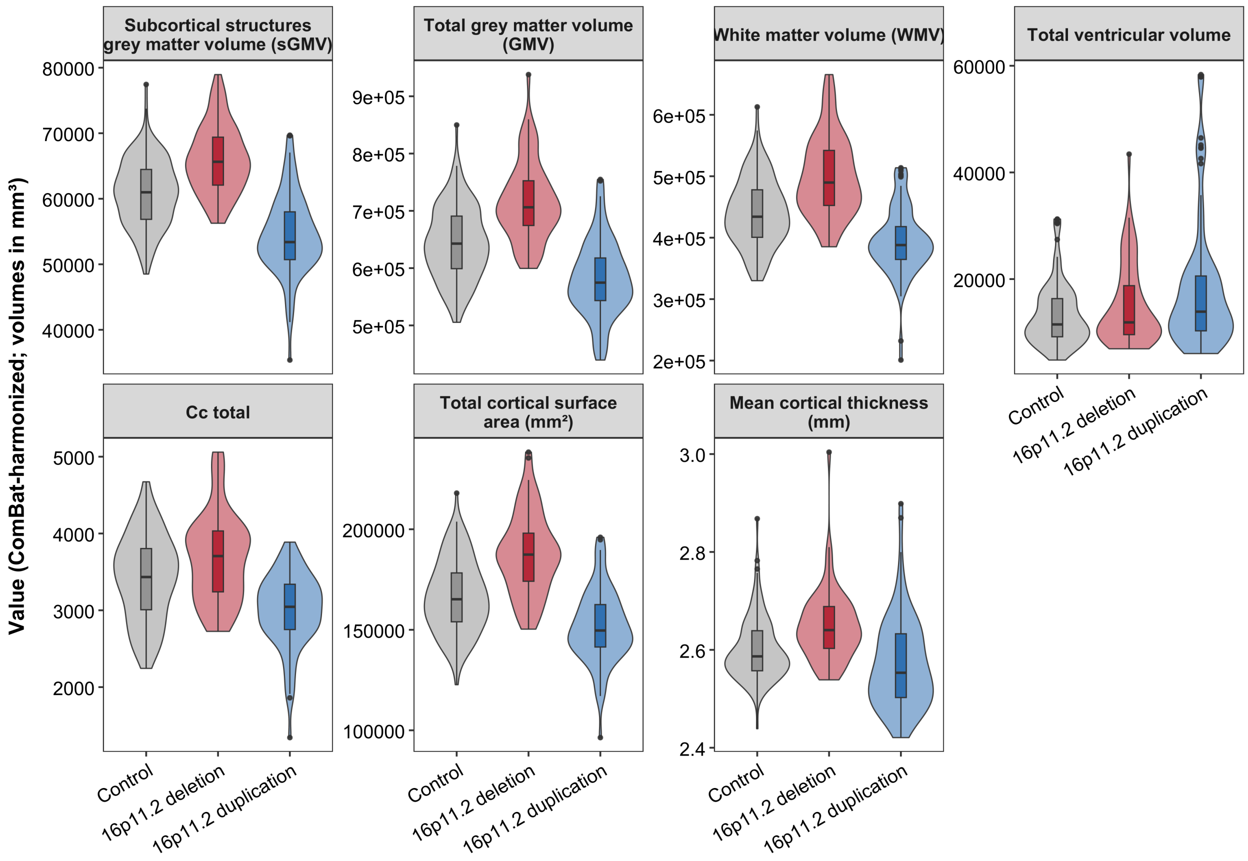


**Supplementary Figure S2: Raw neuroanatomical measure distributions by genotype.** Distributions of unadjusted, ComBat-harmonized neuroanatomical measures are shown for controls, deletion, and duplication across cortical regions, subcortical and other aseg-derived measures, and global/tissue-compartment measures. Violin plots show the observed distributions, with embedded box plots indicating the median and interquartile range.

#### Figure S3: Comparison of Brain-Size Adjustment Method

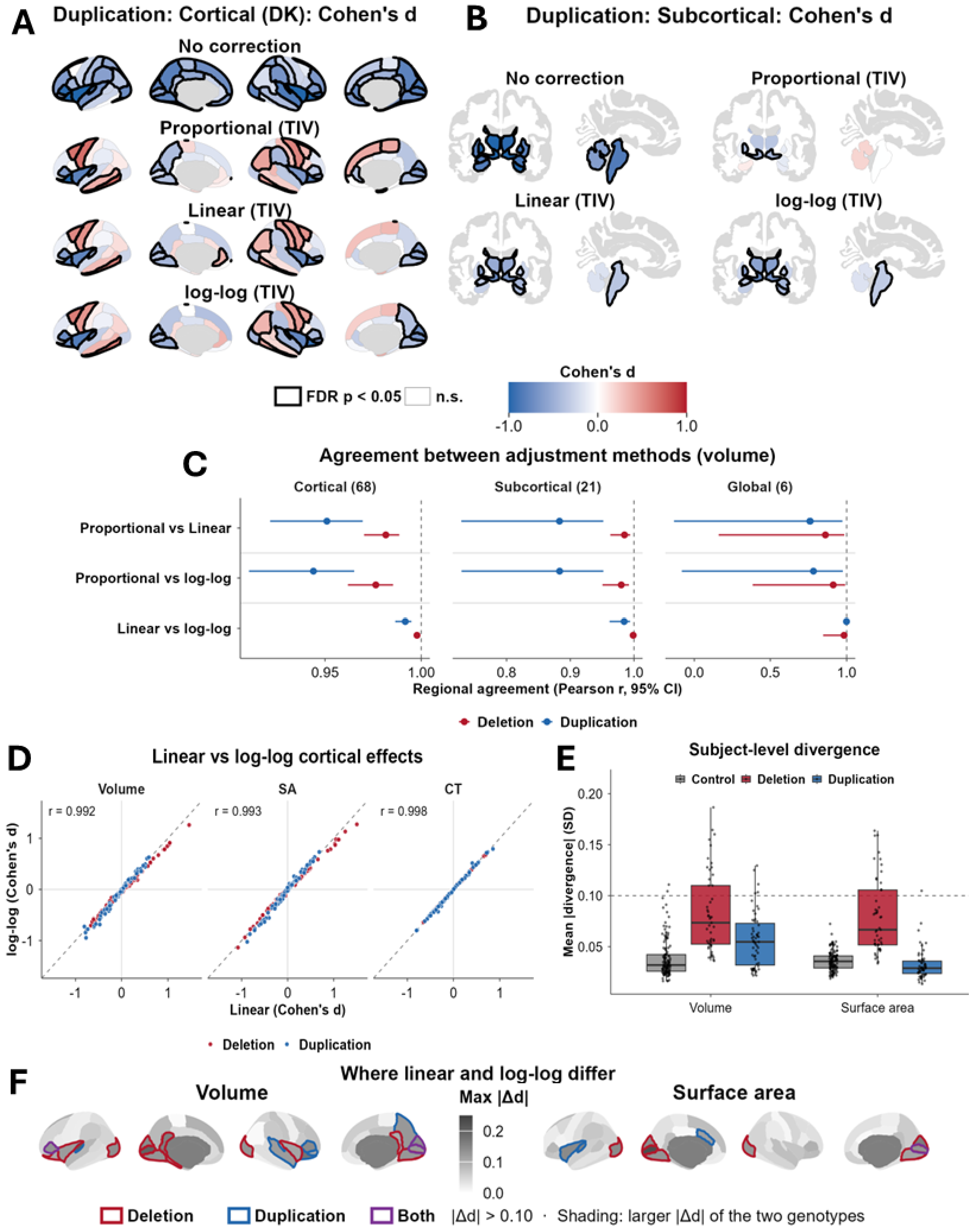


**Supplementary Figure S3: Comparison of brain-size adjustment methods.** (A) Cortical and (B) subcortical regional volume effects (Cohen's d) in duplication under no correction, proportional normalization, linear TIV adjustment, and log-log TIV adjustment; black outlines indicate FDR $p < 0.05$. (C) Pairwise agreement between methods in regional Cohen's d, by anatomical class and genotype (Pearson r with 95% CI). (D) Linear versus log-log cortical effects for volume, SA, and CT; the dashed line indicates identity. (E) Subject-level mean absolute divergence between linear and log-log predictions across the regions outlined in (F), in regional SD units; the dashed line marks 0.10. (F) Absolute difference between linear and log-log Cohen's d (|Δd|) for volume and SA, shaded by the larger of the two genotypes; outlines mark |Δd| > 0.10 in deletion (red), duplication (blue), or both (purple).

#### Figure S4: Sensitivity to Global Reference Measure

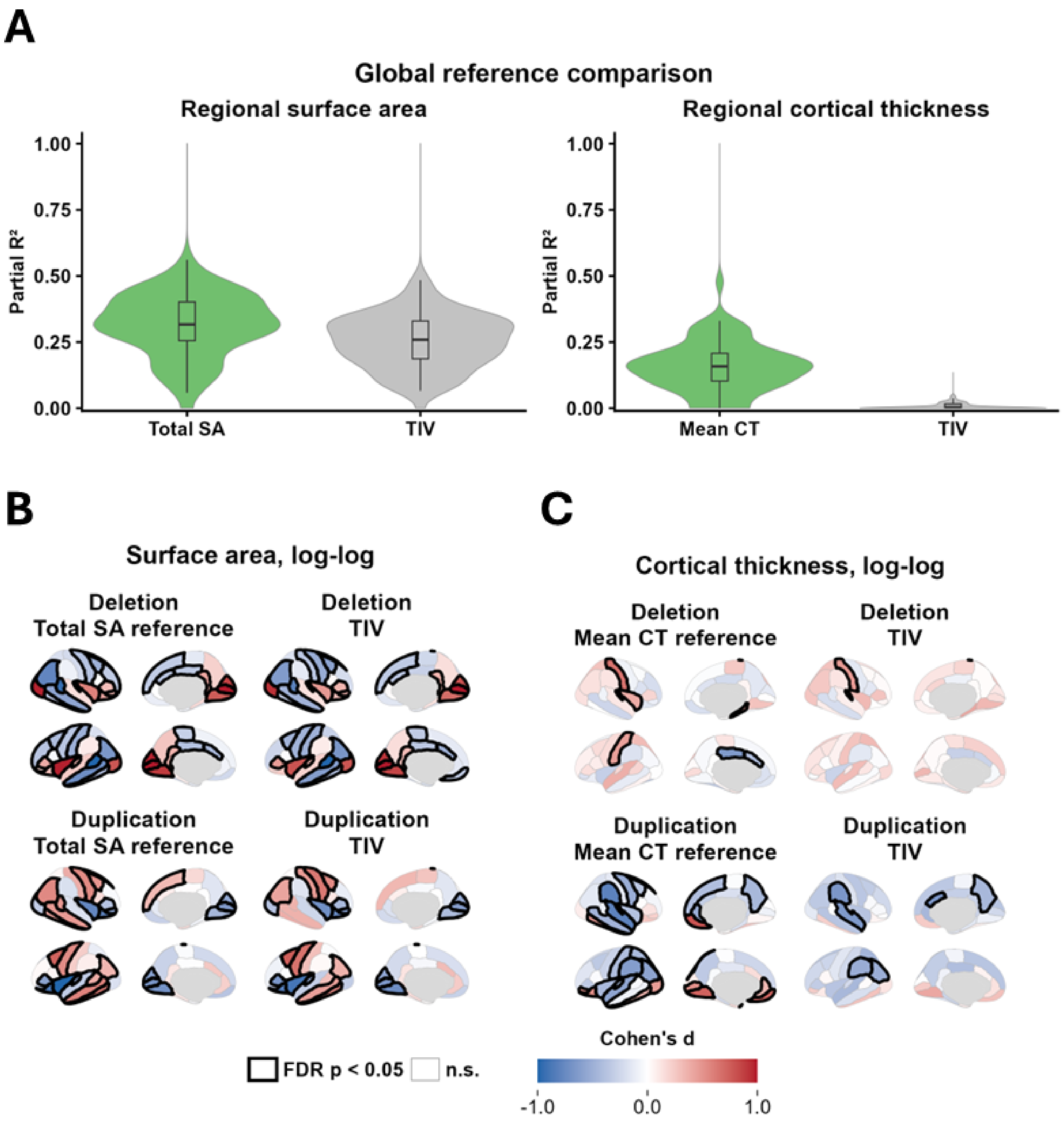


**Supplementary Figure S4: Sensitivity of cortical disproportionate effects to the global reference measure.** Supplementary Figure S4: Sensitivity of cortical disproportionate effects to the global reference measure. Regional SA and CT were adjusted for overall size using each metric's own global measure (total SA or mean CT) rather than TIV. (A) Variance in regional SA and CT explained by each candidate reference in controls. For each of the 68 cortical regions, partial $R^2$ is the proportion of variance explained by the reference measure after adjustment for age (spline), sex, and mean bilateral Euler number; violins show distributions across regions. Regional SA was explained similarly by total SA and TIV, whereas regional CT was explained by mean CT but was essentially unrelated to TIV. (B) Regional SA effects and (C) regional CT effects from the log-log model, re-estimated with TIV as the global reference and shown alongside the primary model for deletion and duplication. SA effects were similar with either reference. CT effects retained a similar spatial pattern with TIV as the reference, but fewer regions survived correction. Black outlines indicate FDR $p < 0.05$.

#### Figure S5: Volume spatial correspondence and dosage-response patterns before TIV adjustment

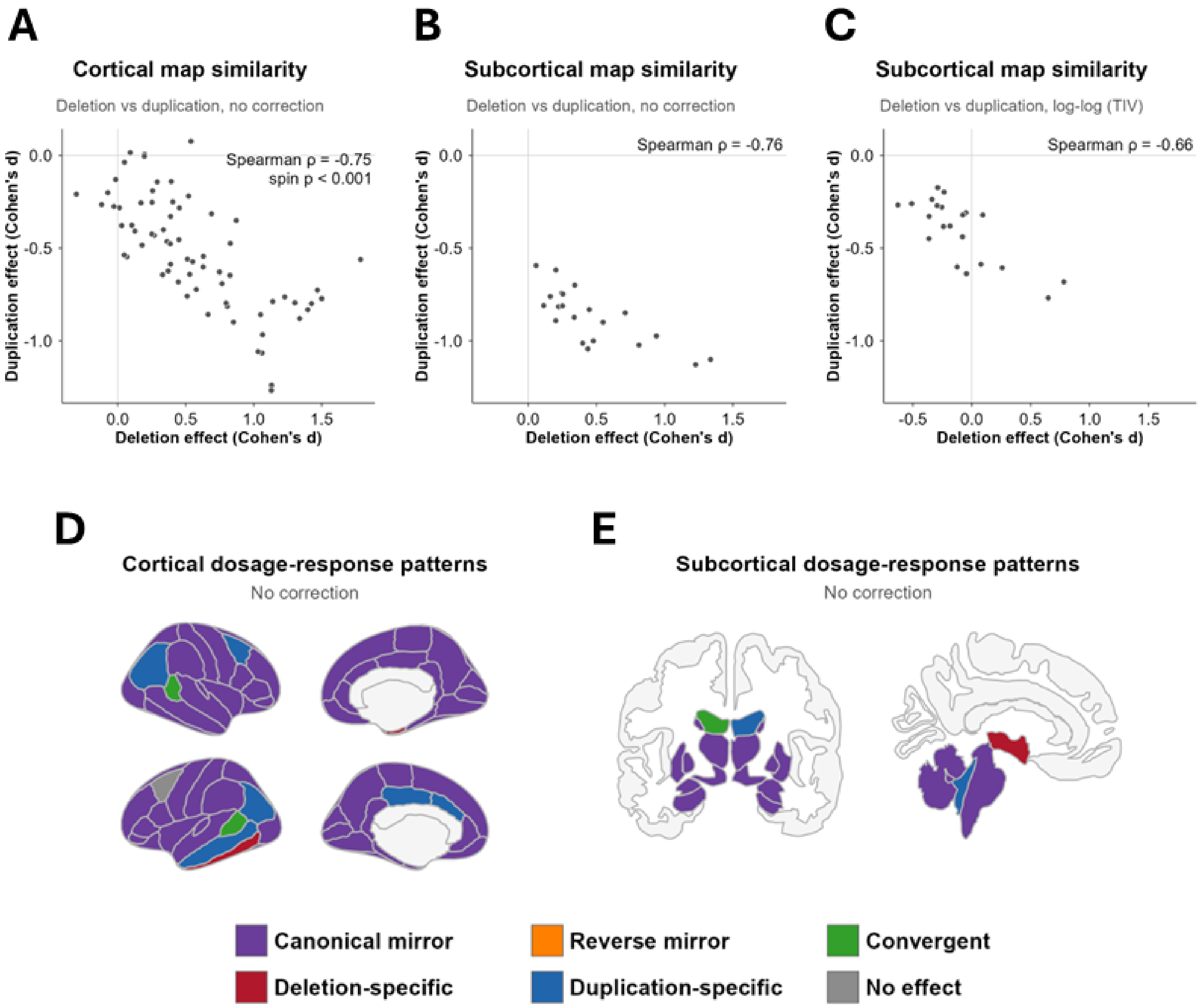


**Supplementary Figure S5: Spatial correspondence and dosage-response patterns of regional volume before TIV adjustment, with subcortical correspondence after adjustment.** (A) Relationship between TIV-unadjusted cortical effects in deletion and duplication. Spearman's ρ summarizes rank correspondence between regional effect maps, with two-sided significance assessed using a spherical spin test preserving cortical spatial autocorrelation. (B) Relationship between TIV-unadjusted subcortical effects in deletion and duplication, summarized by Spearman's ρ without a significance test. (C) The same subcortical relationship after log-log TIV adjustment, also summarized by Spearman's ρ without a significance test. (D) TIV-unadjusted cortical dosage-response patterns classified using the framework defined in Fig. 1E. (E) TIV-unadjusted aseg-derived dosage-response patterns classified using the same framework.

#### Figure S6: TIV-unadjusted spatial correspondence and dosage-response patterns in SA and CT

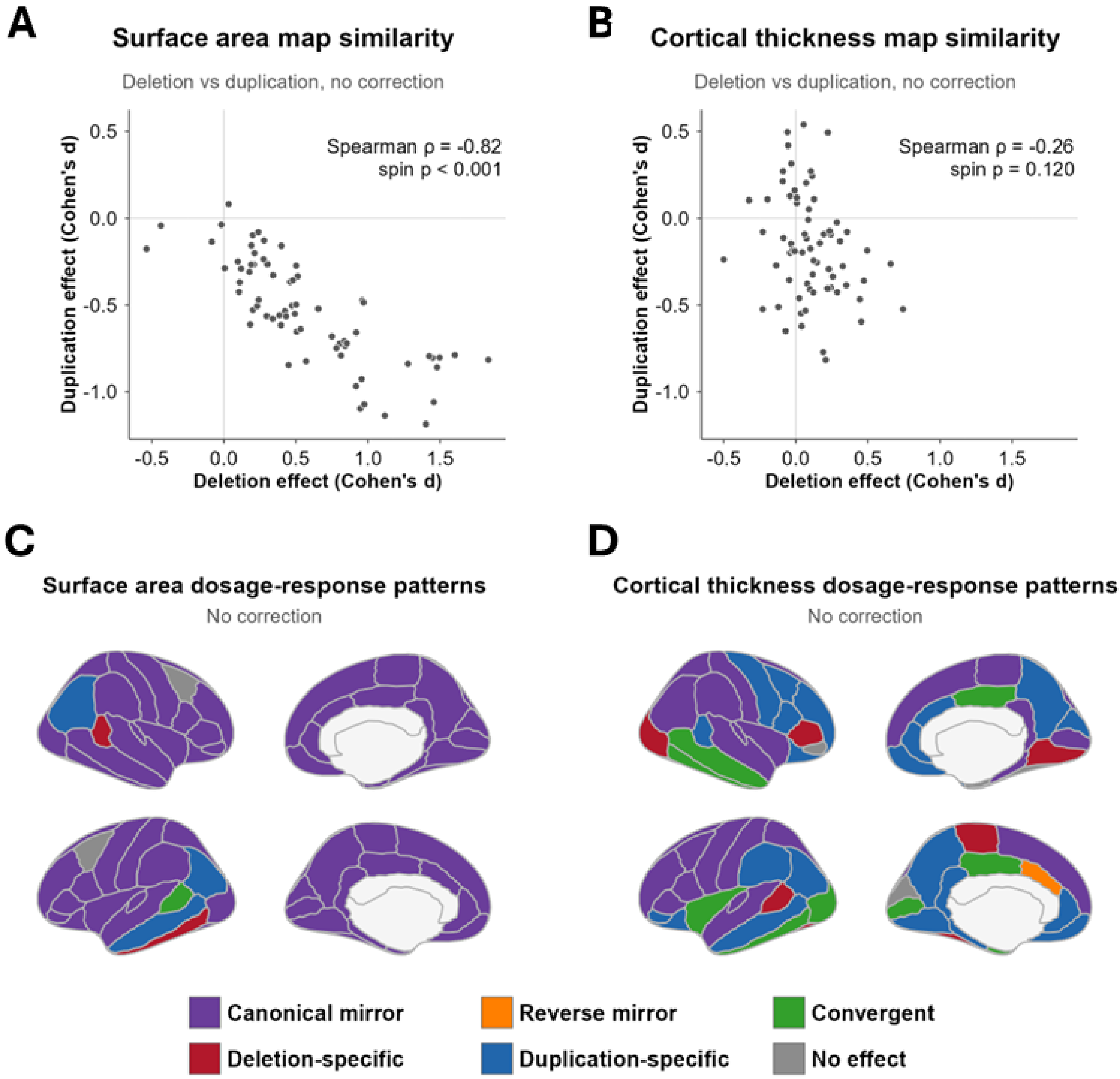


**Supplementary Figure S6: TIV-unadjusted spatial correspondence and dosage-response patterns in cortical surface area and thickness.** (A) Relationship between TIV-unadjusted cortical surface area effects in deletion and duplication. Spearman's ρ summarizes rank correspondence between regional effect maps, with two-sided significance assessed using a spherical spin test preserving cortical spatial autocorrelation. (B) Relationship between TIV-unadjusted cortical thickness effects in deletion and duplication, with significance assessed using the same approach. (C) TIV-unadjusted cortical surface area dosage-response patterns classified using the framework defined in Fig. 1E. (D) TIV-unadjusted cortical thickness dosage-response patterns classified using the same framework.

#### Figure S7: Allometric Phenotypes in SA and CT

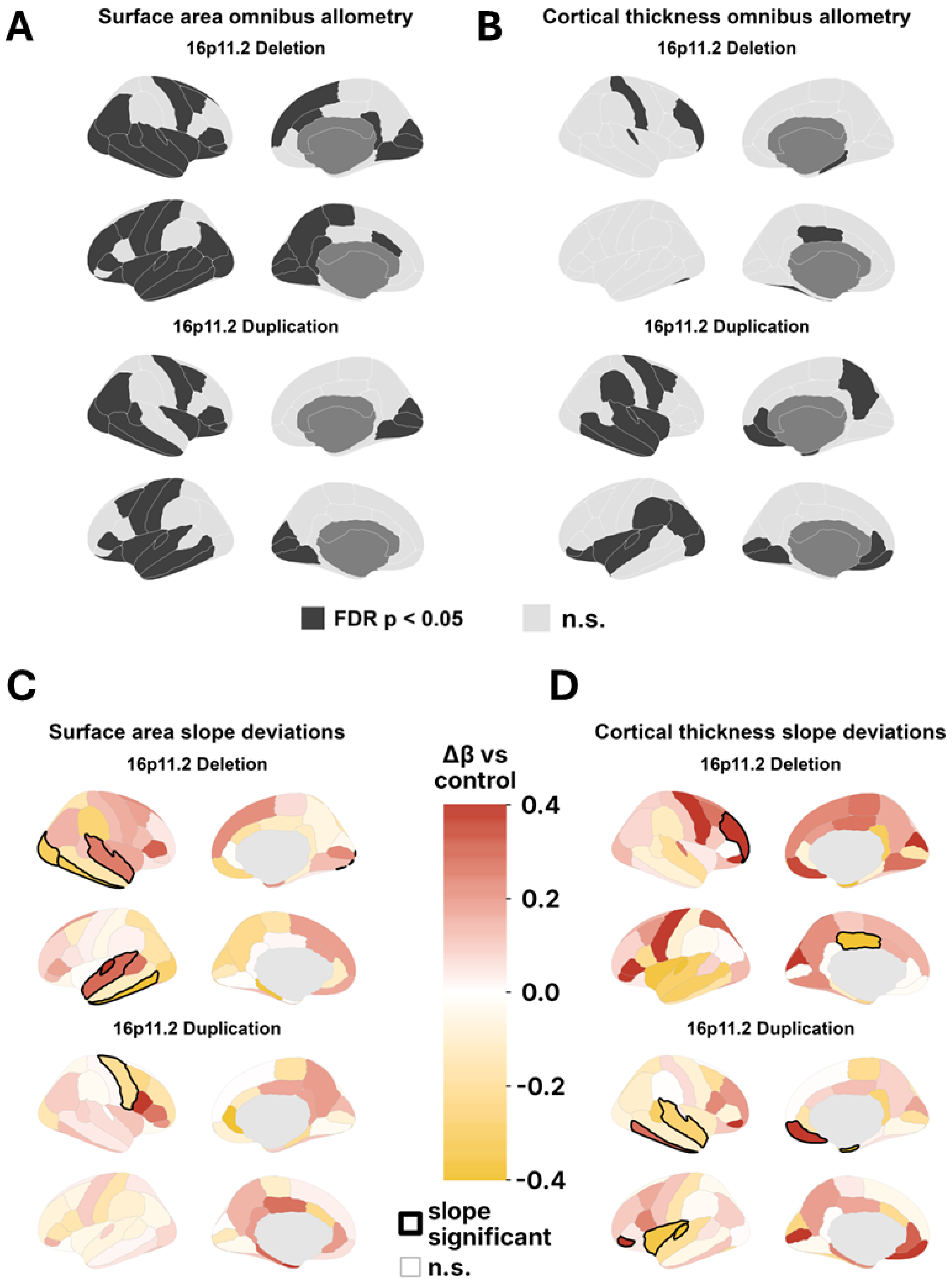


**Supplementary Figure S7: Genotype-specific alterations in cortical surface area and cortical thickness scaling.** (A) SA and (B) CT regions showing genotype-related differences in the allometric scaling relationship, shown separately for deletion and duplication. The omnibus test is conducted separately for each CNV versus controls and detects differences in scaling constant, slope, or both; dark regions survive FDR correction (FDR $p < 0.05$). (C) SA and (D) CT genotype-specific differences in scaling slope ($\Delta\beta$) relative to controls. Positive and negative values indicate steeper and shallower scaling, respectively; black outlines identify omnibus-positive regions with nominally significant slope differences ($p < 0.05$).

### Figure S8: Masking of Cortical Regions with Altered Brain-Size Scaling

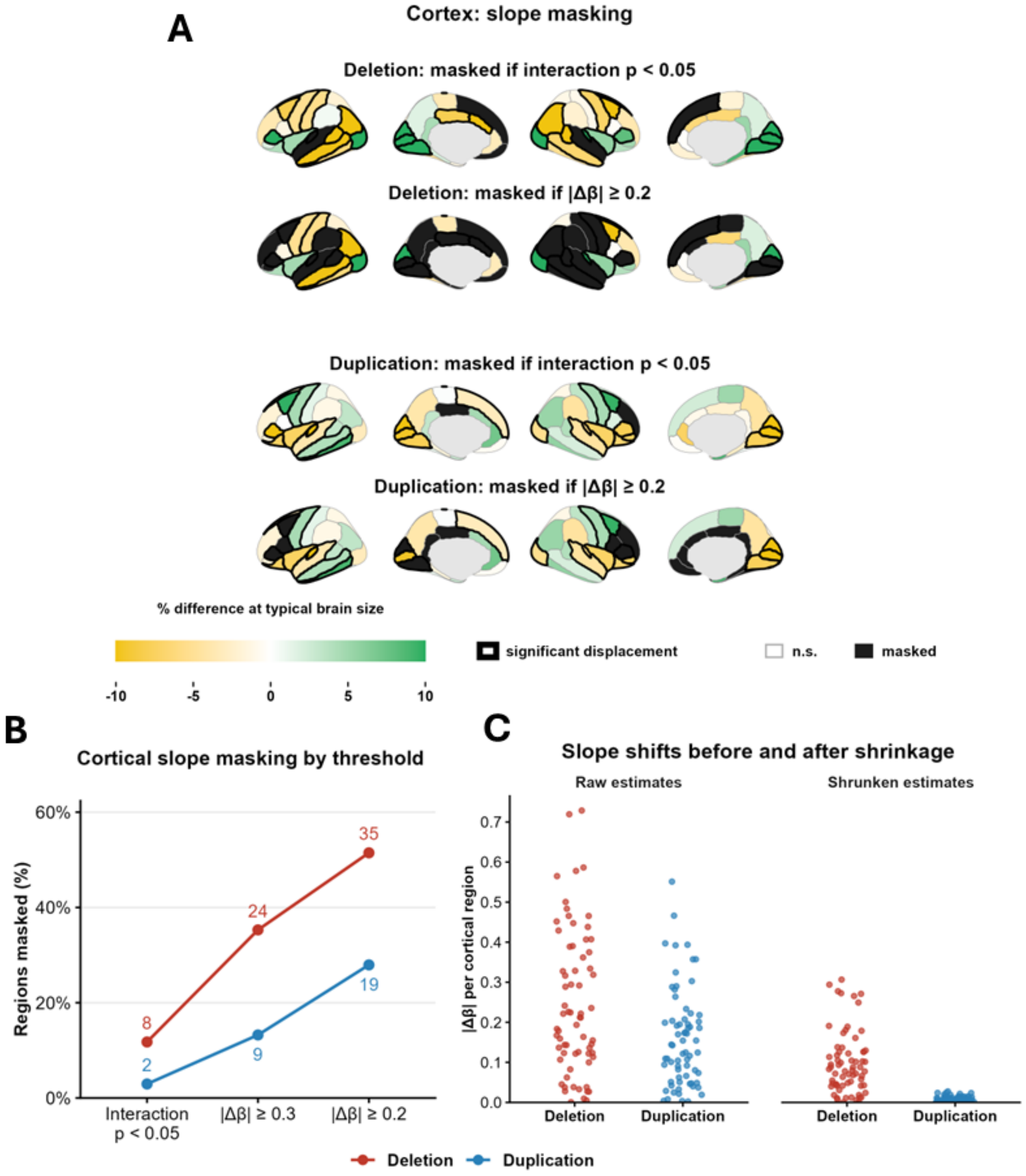


**D**

Subcortical structures masked (of 21)

| CNV | Interaction p < 0.05 | \|Δβ\| ≥ 0.3 | \|Δβ\| ≥ 0.2 | \|Δβ\| ≥ 0.2 after shrinkage |
|---|---|---|---|---|
| Deletion | 2 | 2 | 4 | 0 |
| Duplication | 0 | 0 | 3 | 0 |

All significantly disproportionate subcortical structures (7 in deletion, 13 in duplication) remained unmasked under every rule.

**Supplementary Figure S8: Masking of cortical regions with altered brain-size scaling.** (A) Estimated percentage difference from controls in regional cortical volume at the typical unrelated-control brain size, for deletion (top two rows) and duplication (bottom two rows). In each CNV, the first row masks omnibus-positive regions with a nominally significant genotype-by-size interaction ($p < 0.05$), and the second masks regions whose slope difference reaches the 5% tolerance ($|\Delta\beta| \geq 0.2$; Supplementary Methods), regardless of significance. Masked regions are shown in black; outlines identify nominally significant differences in the scaling constant. (B) Percentage of the 68 cortical regions masked under interaction-significance masking and under the 10% ($|\Delta\beta| \geq 0.3$) and 5% ($|\Delta\beta| \geq 0.2$) tolerances; numbers give region counts. (C) Absolute slope differences ($|\Delta\beta|$) for each cortical region before (raw estimates) and after empirical Bayes shrinkage toward zero (shrunken estimates; Supplementary Methods). (D) Number of the 21 subcortical structures masked under each rule, including the 5% tolerance applied after shrinkage; all significantly disproportionate subcortical structures remained unmasked under every rule.

### Supplementary Figure S9. Regional allometric fits — cortical (1/12)

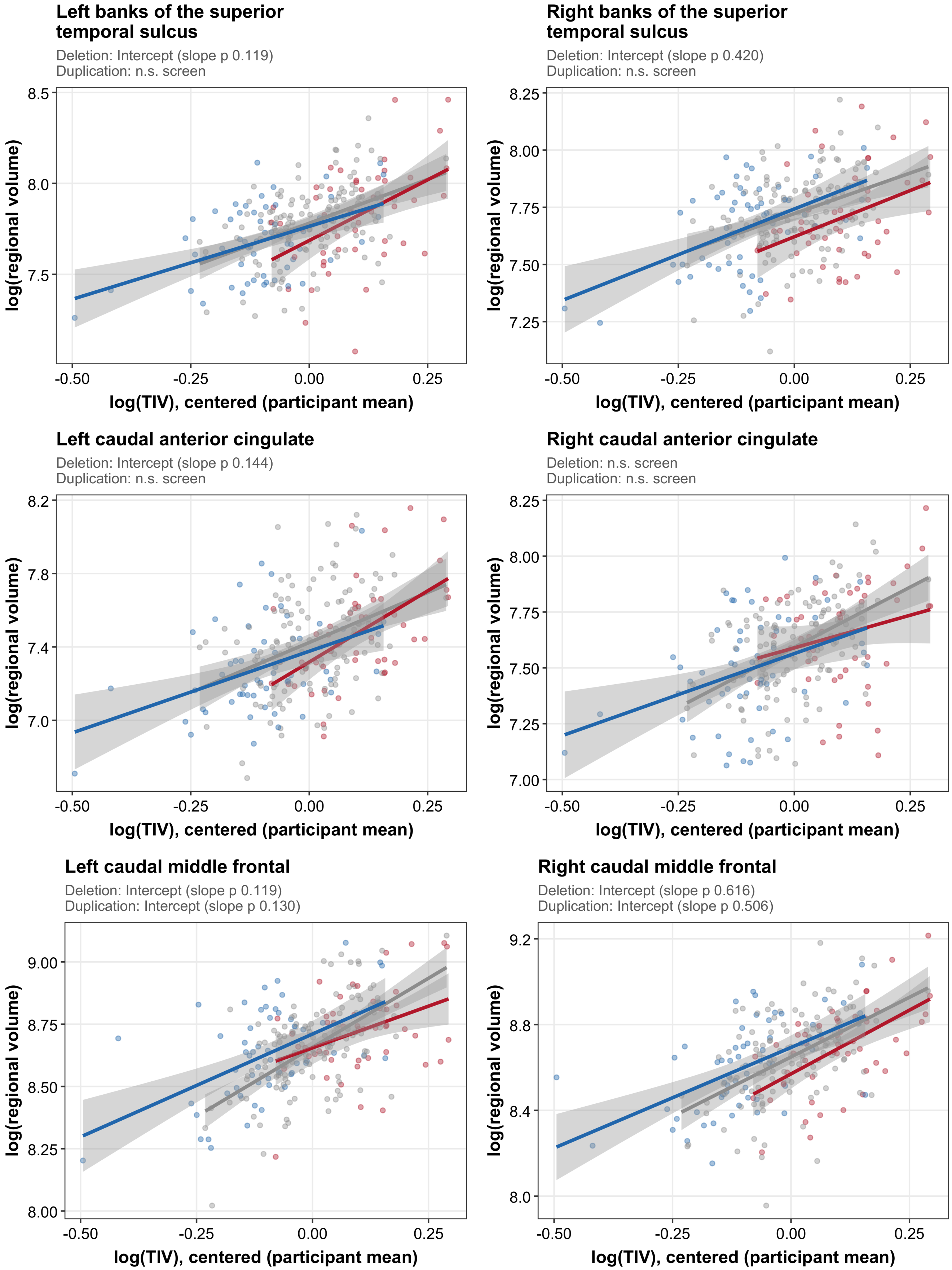

#### Supplementary Figure S9. Regional allometric fits — cortical (2/12)

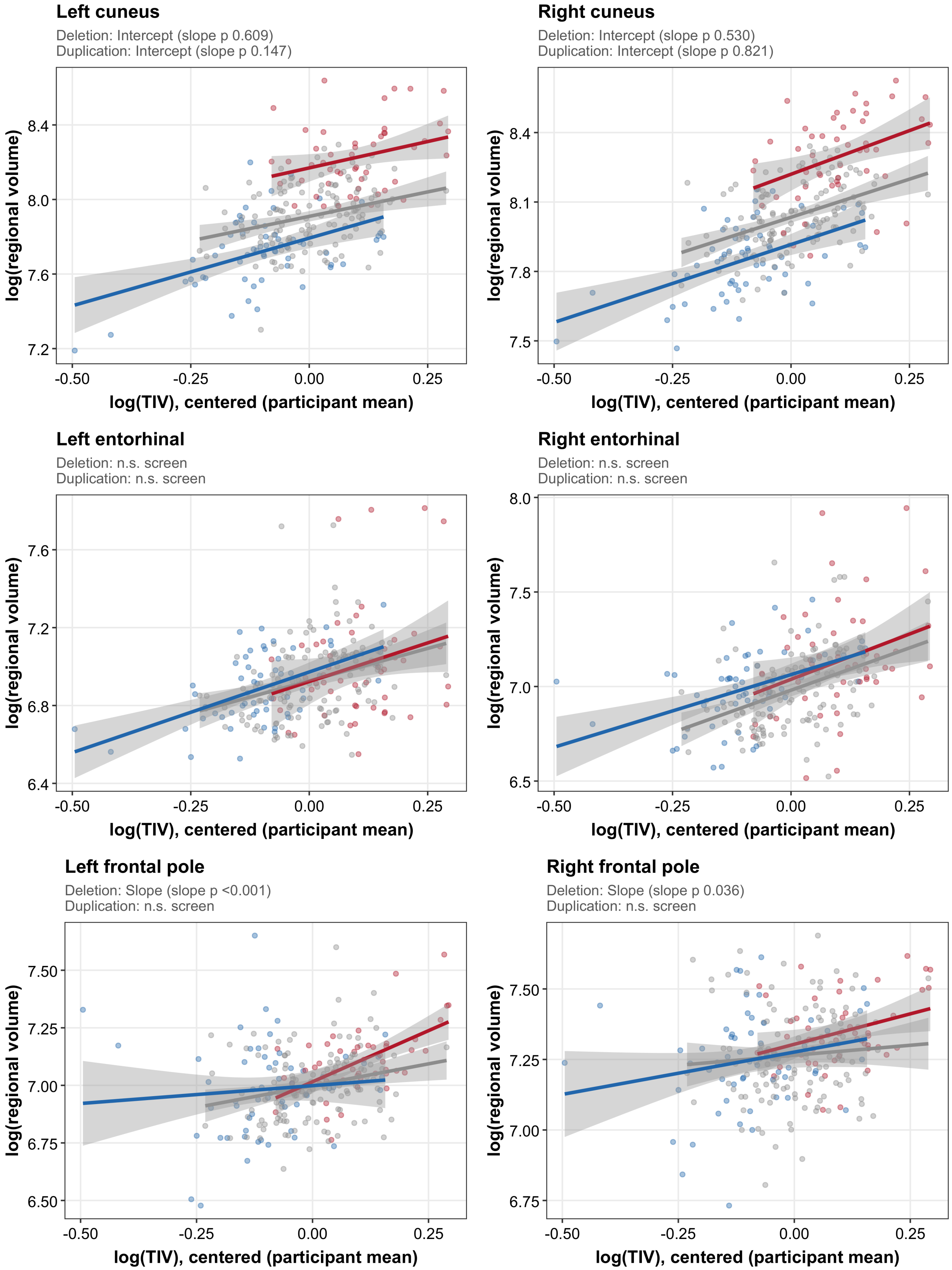

### Supplementary Figure S9. Regional allometric fits — cortical (3/12)

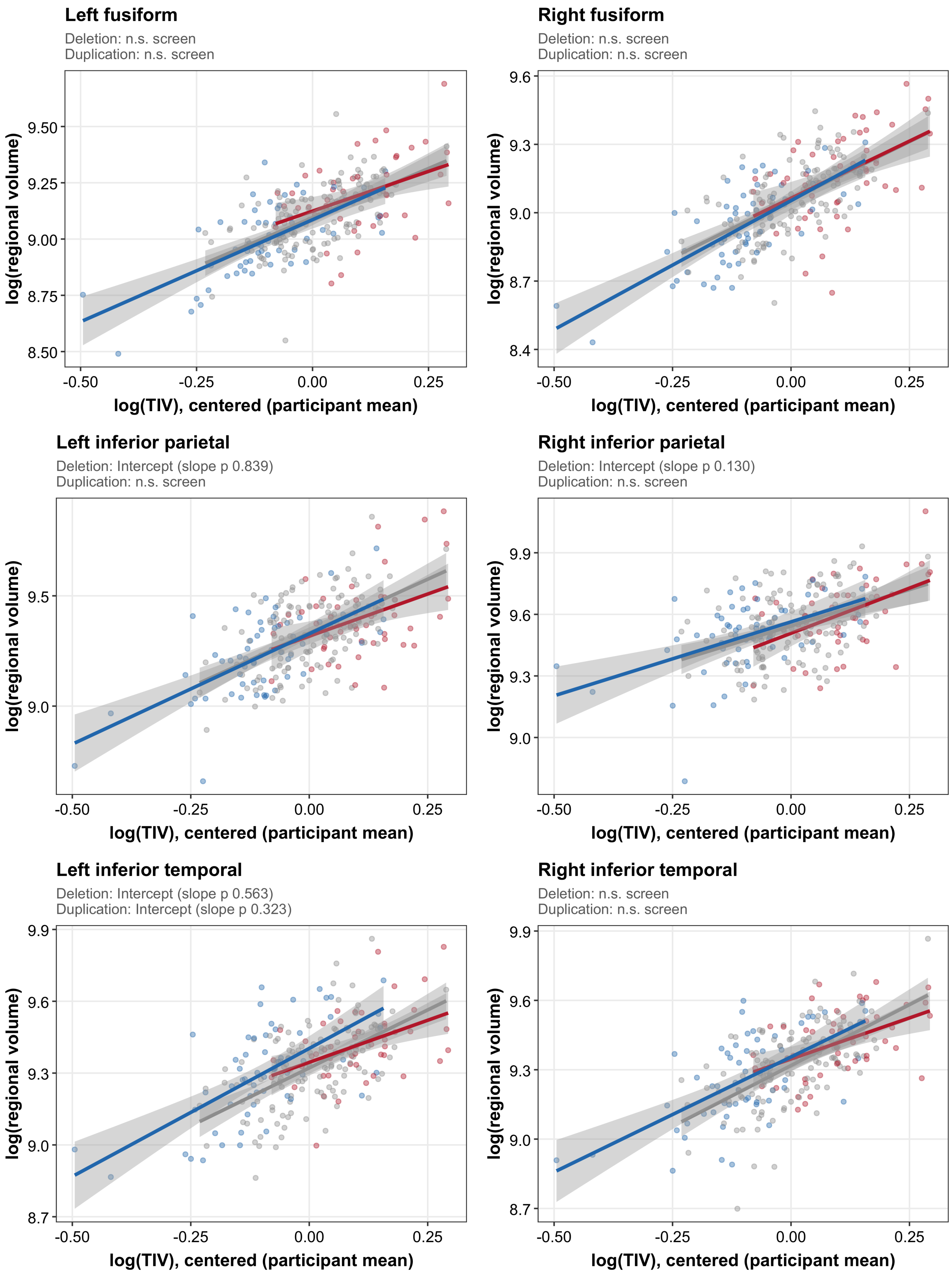

### Supplementary Figure S9. Regional allometric fits — cortical (4/12)

##### Left insula

Deletion: Intercept (slope p 0.285)
Duplication: Intercept (slope p 0.940)

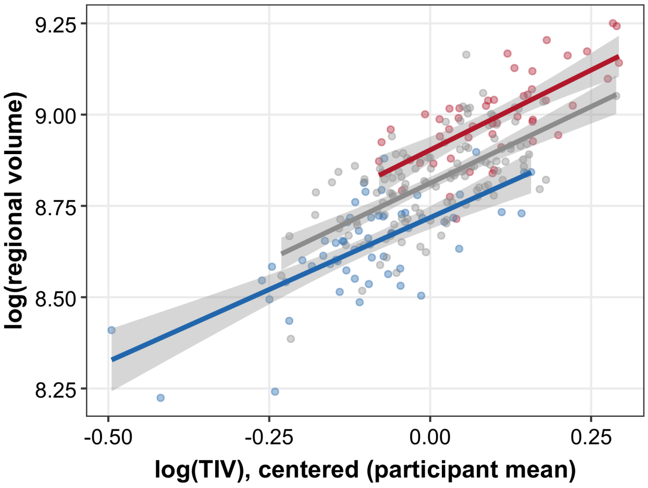


##### Right insula

Deletion: Intercept (slope p 0.950)
Duplication: Intercept (slope p 0.442)

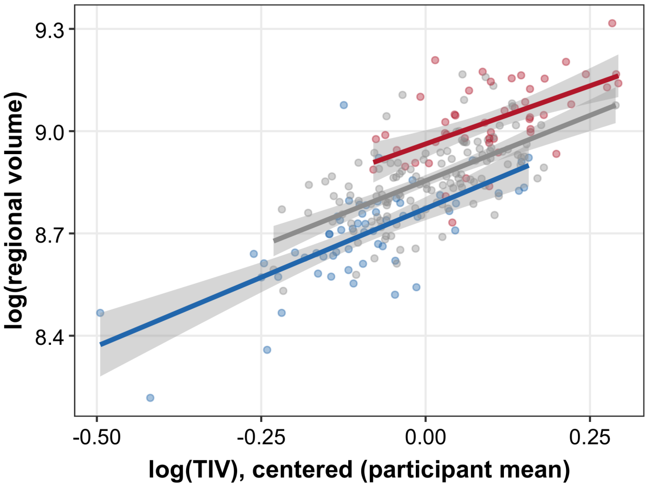


##### Left isthmus cingulate

Deletion: n.s. screen
Duplication: n.s. screen

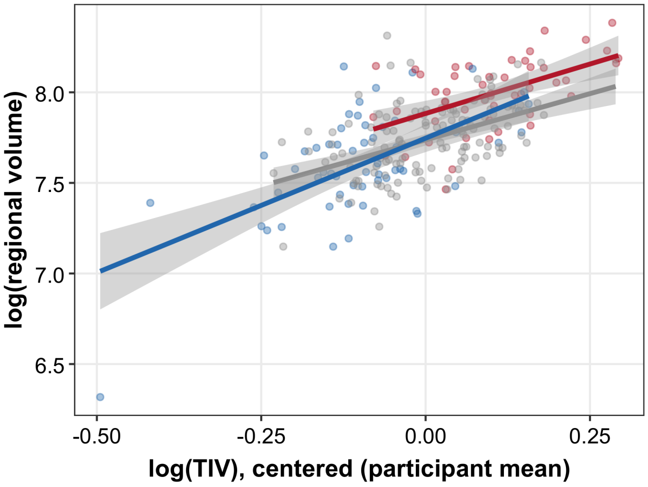


##### Right isthmus cingulate

Deletion: n.s. screen
Duplication: n.s. screen

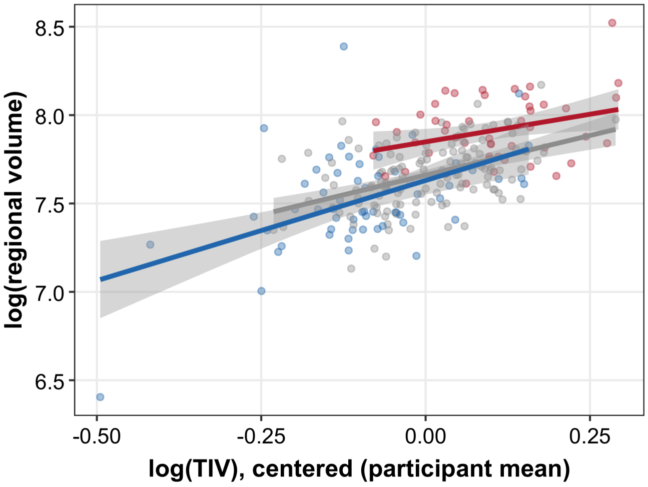


##### Left lateral occipital

Deletion: Intercept (slope p 0.365)
Duplication: n.s. screen

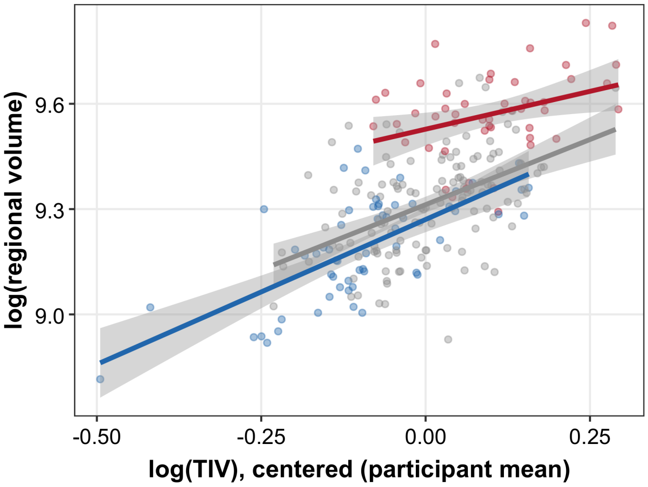


##### Right lateral occipital

Deletion: Intercept (slope p 0.891)
Duplication: Intercept (slope p 0.510)

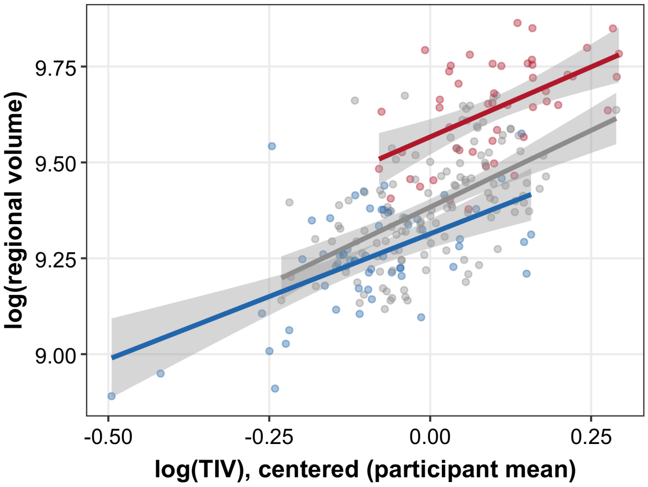

### Supplementary Figure S9. Regional allometric fits — cortical (5/12)

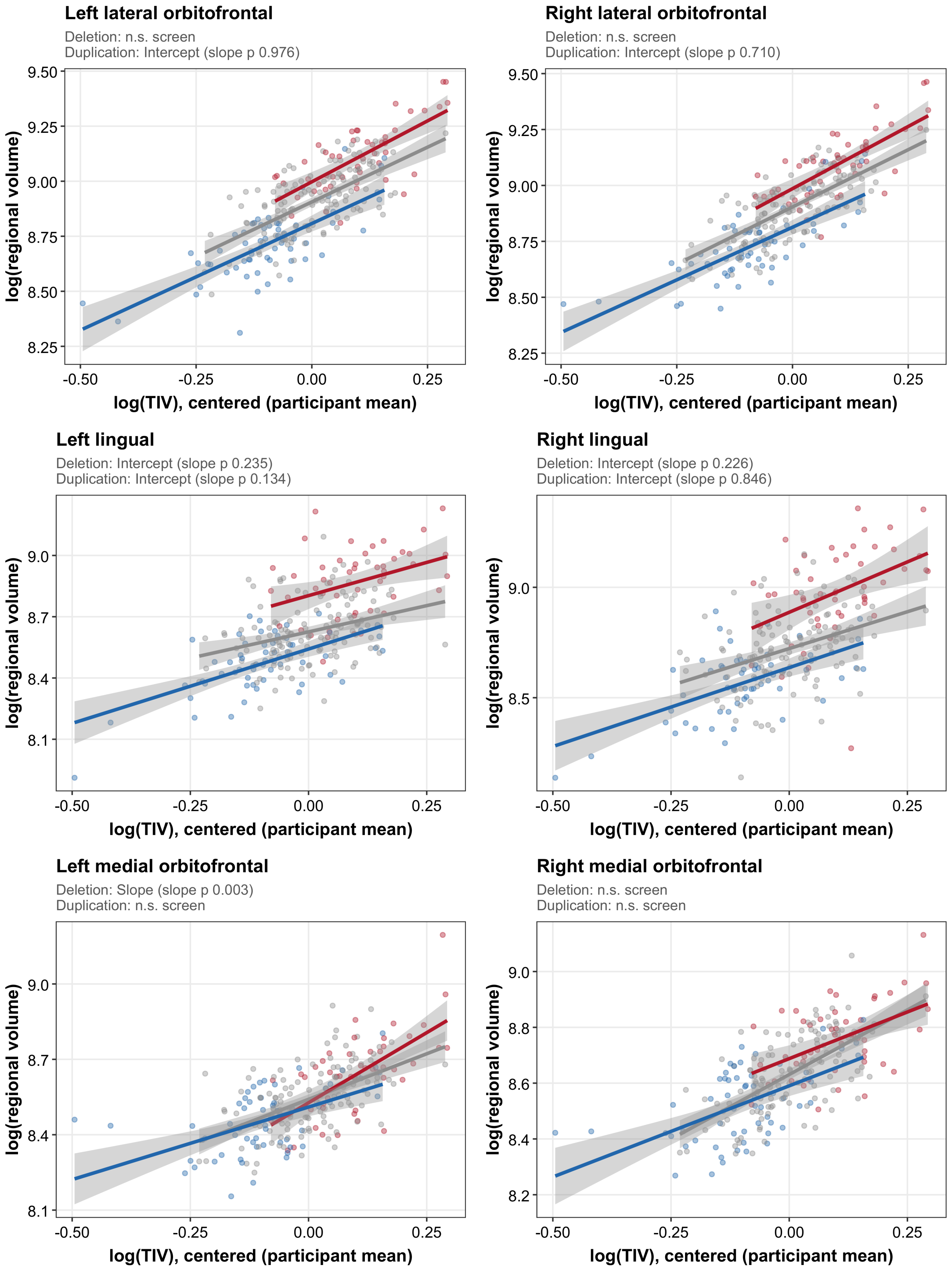

### Supplementary Figure S9. Regional allometric fits — cortical (6/12)

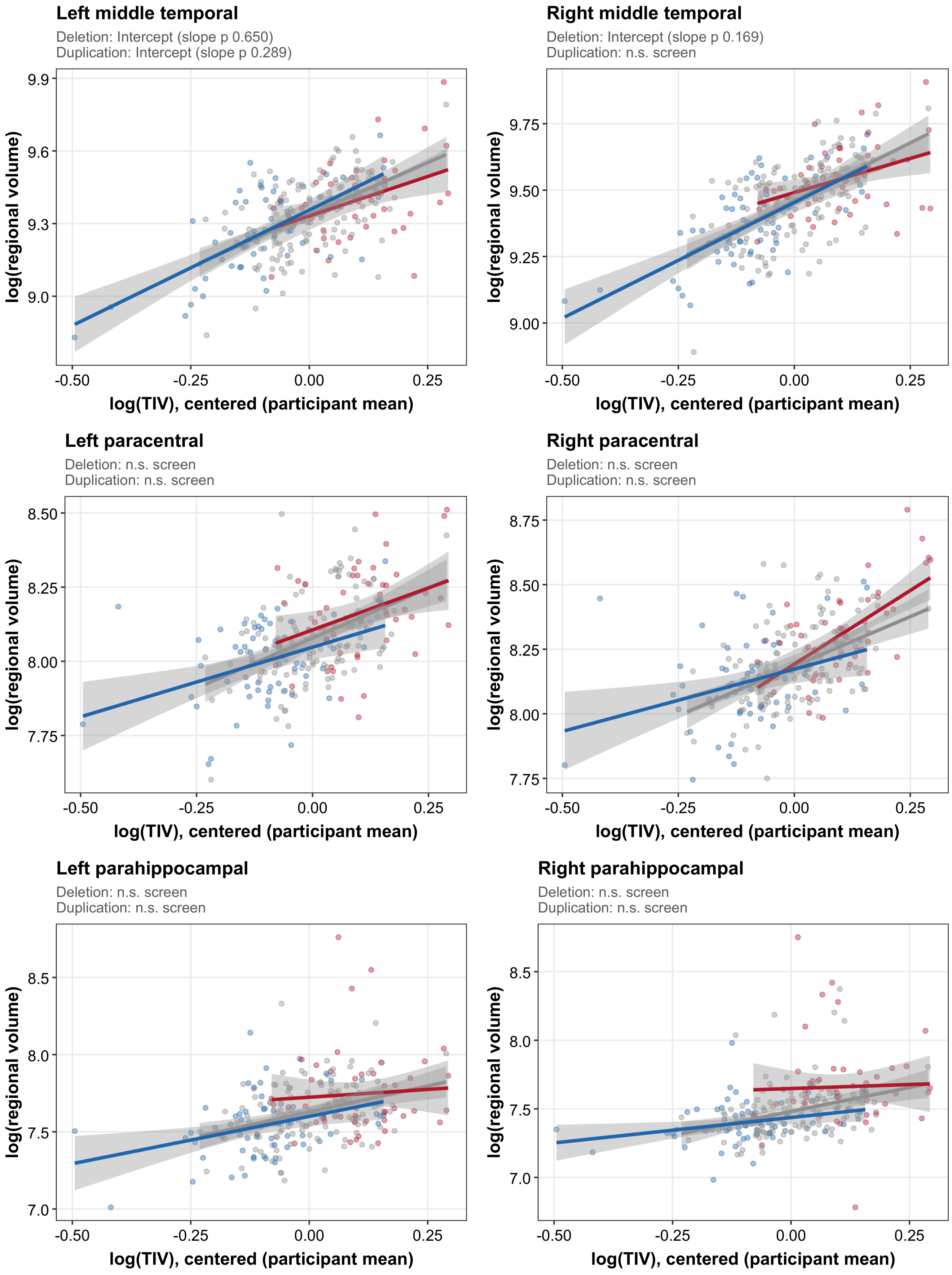

### Supplementary Figure S9. Regional allometric fits — cortical (7/12)

**Left pars opercularis**

Deletion: n.s. screen
Duplication: n.s. screen

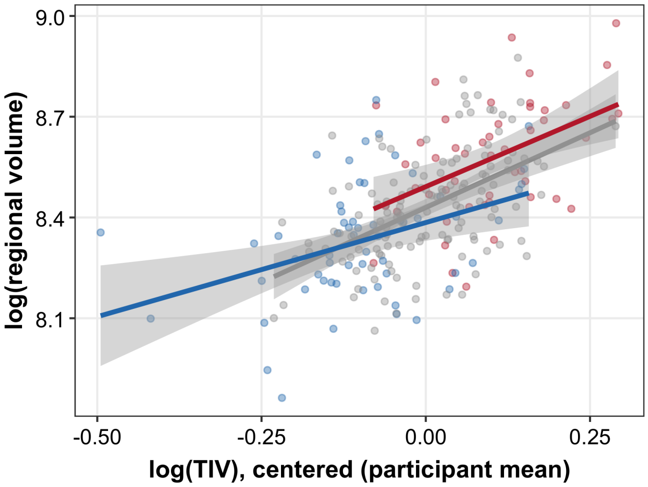


**Right pars opercularis**

Deletion: n.s. screen
Duplication: n.s. screen

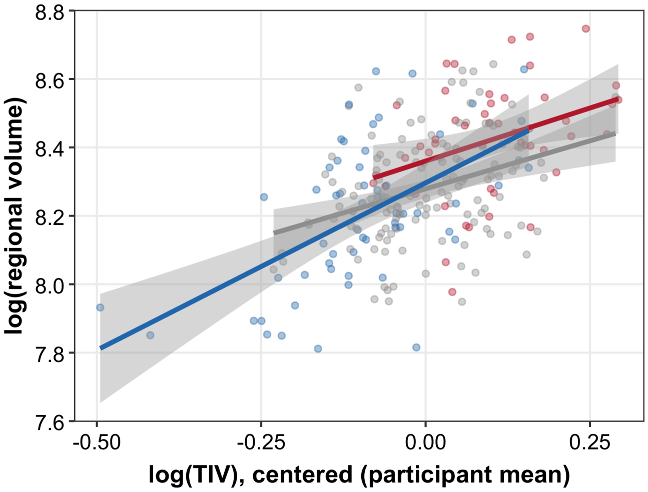


**Left pars orbitalis**

Deletion: n.s. screen
Duplication: n.s. screen

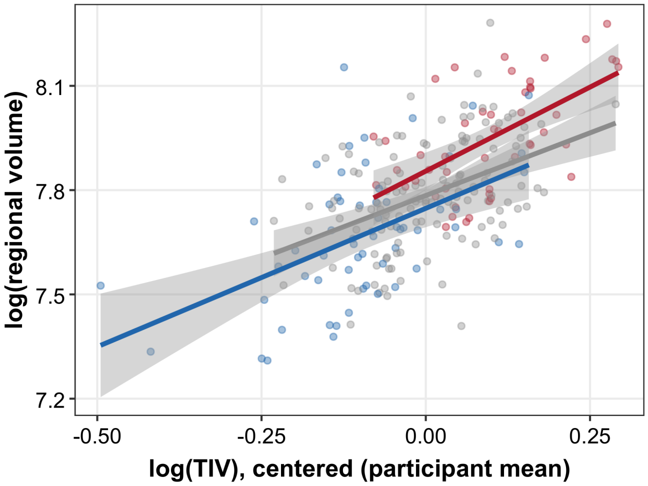


**Right pars orbitalis**

Deletion: n.s. screen
Duplication: Intercept (slope p 0.757)

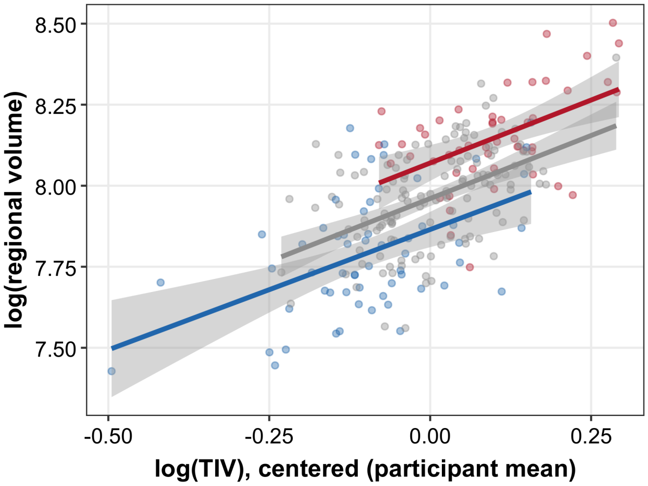


**Left pars triangularis**

Deletion: Intercept (slope p 0.235)
Duplication: Intercept (slope p 0.295)

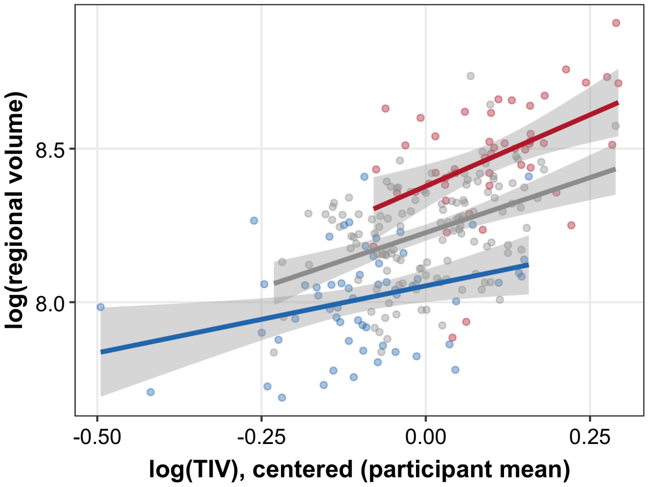


**Right pars triangularis**

Deletion: Intercept (slope p 0.137)
Duplication: Intercept (slope p 0.093)

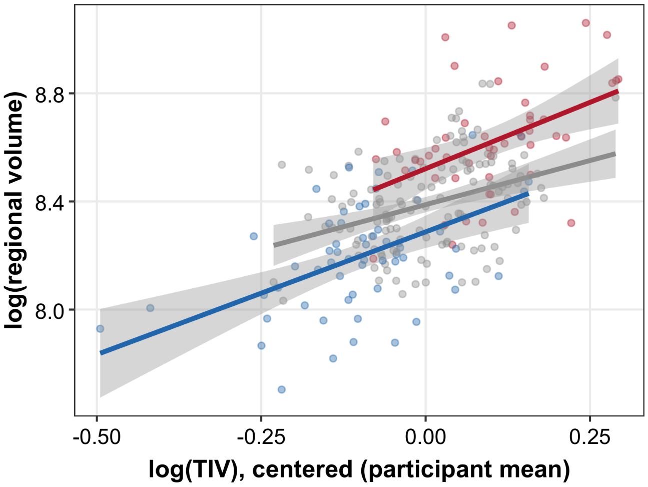

#### Supplementary Figure S9. Regional allometric fits — cortical (8/12)

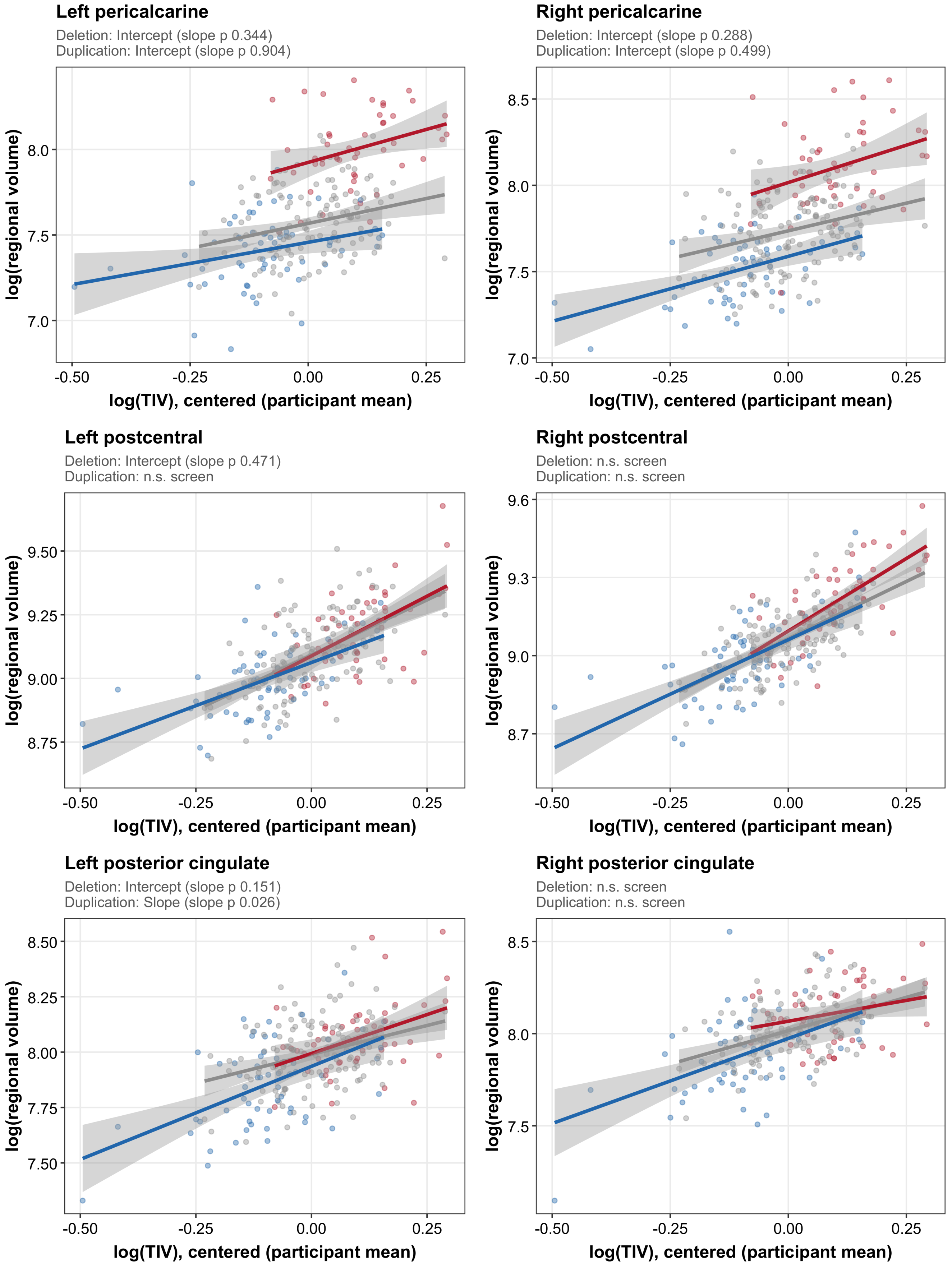

### Supplementary Figure S9. Regional allometric fits — cortical (9/12)

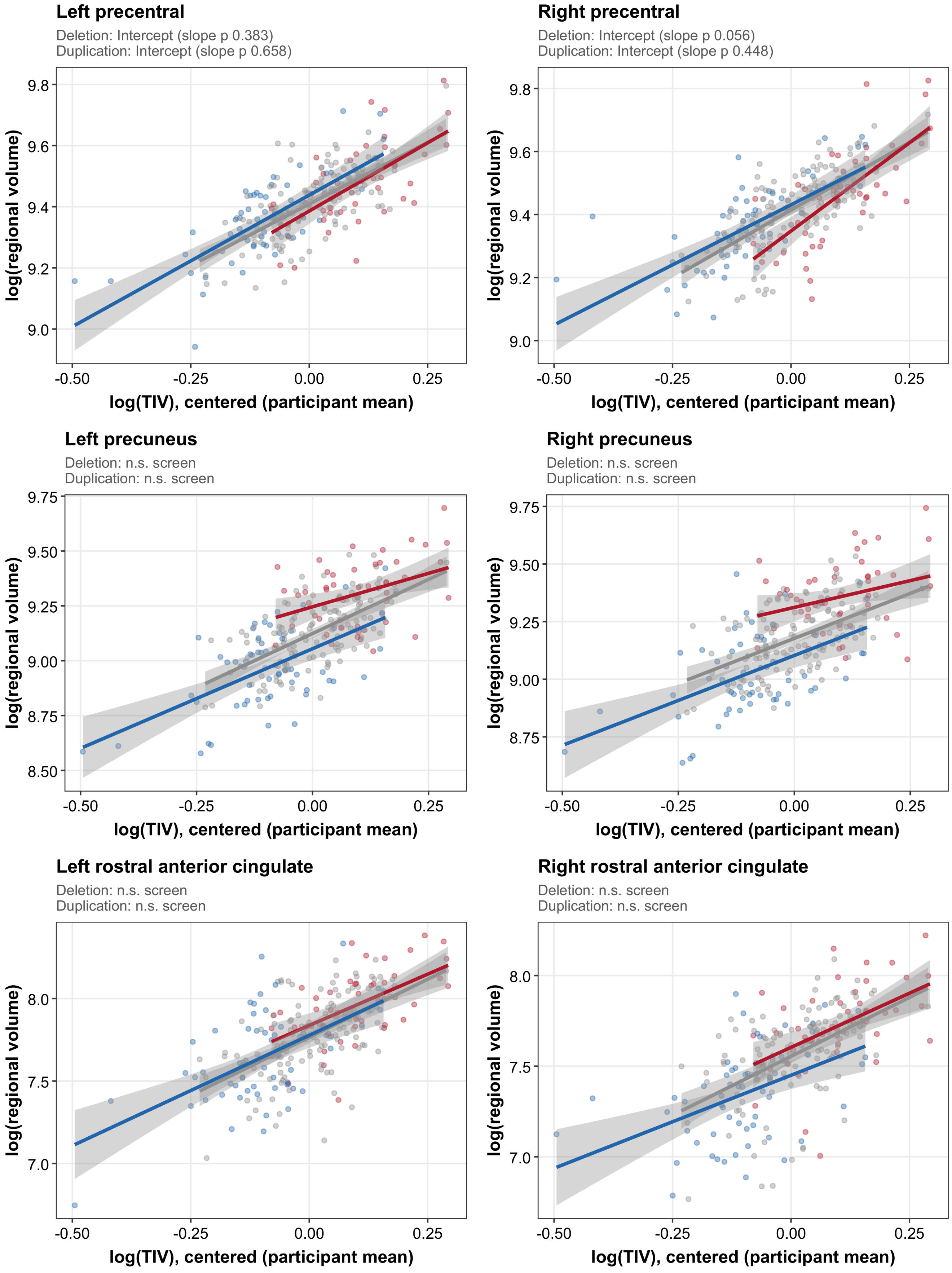

### Supplementary Figure S9. Regional allometric fits — cortical (10/12)

##### Left rostral middle frontal

Deletion: Neither (slope p 0.052)
Duplication: n.s. screen

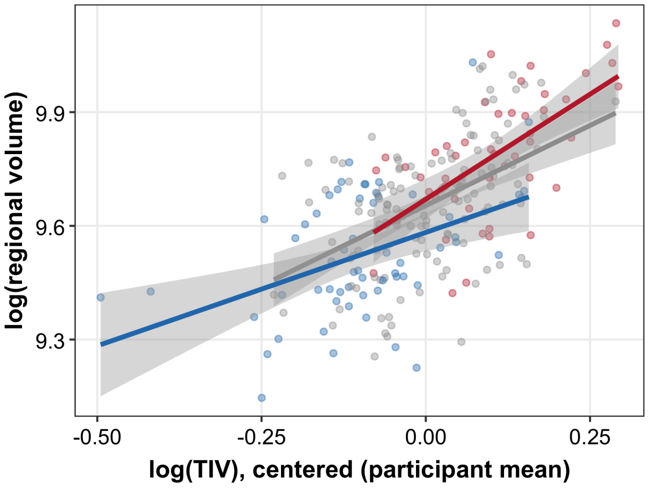


##### Right rostral middle frontal

Deletion: Neither (slope p 0.428)
Duplication: Slope (slope p 0.017)

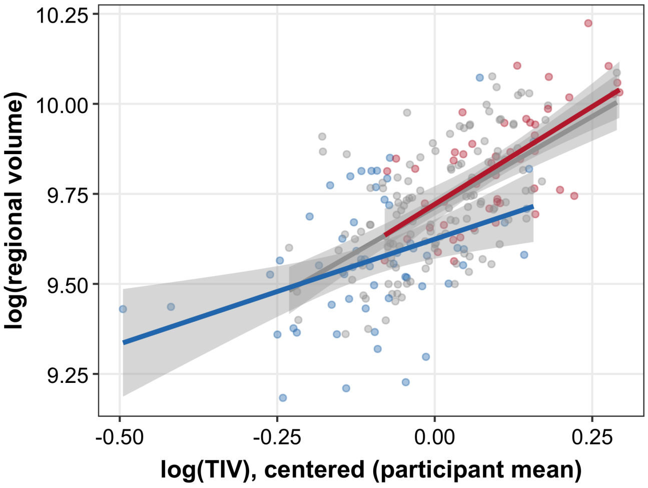


##### Left superior frontal

Deletion: Slope (slope p 0.006)
Duplication: Intercept (slope p 0.985)

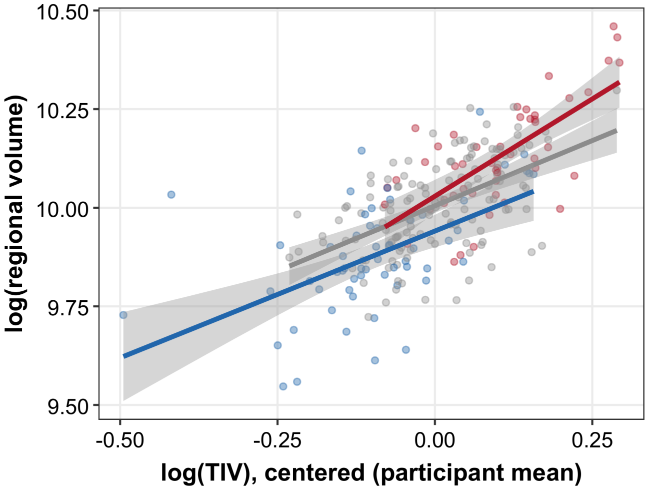


##### Right superior frontal

Deletion: Slope (slope p 0.003)
Duplication: n.s. screen

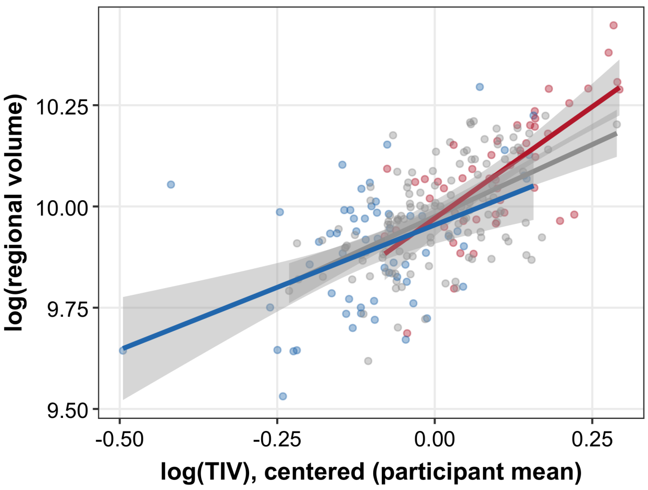


##### Left superior parietal

Deletion: n.s. screen
Duplication: n.s. screen

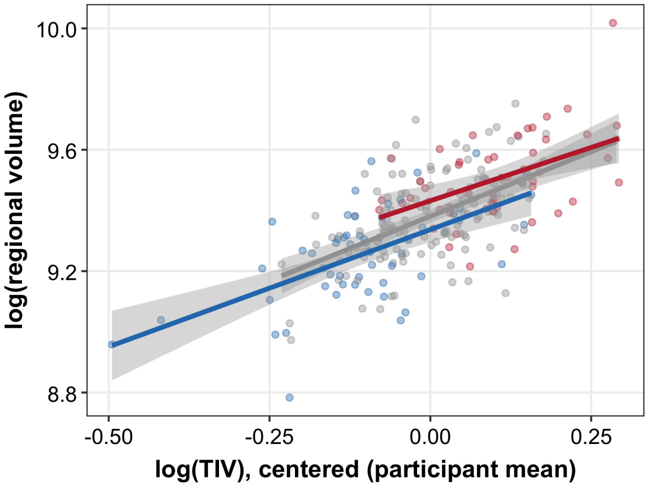


##### Right superior parietal

Deletion: n.s. screen
Duplication: n.s. screen

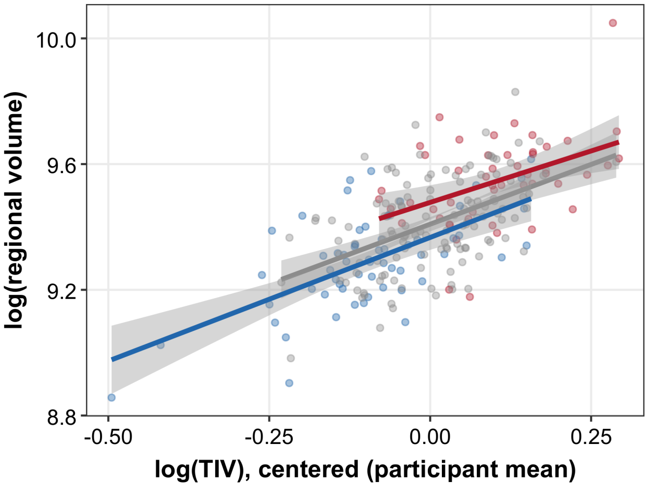

### Supplementary Figure S9. Regional allometric fits — cortical (11/12)

#### Left superior temporal

Deletion: Slope (slope p 0.001)
Duplication: Intercept (slope p 0.556)

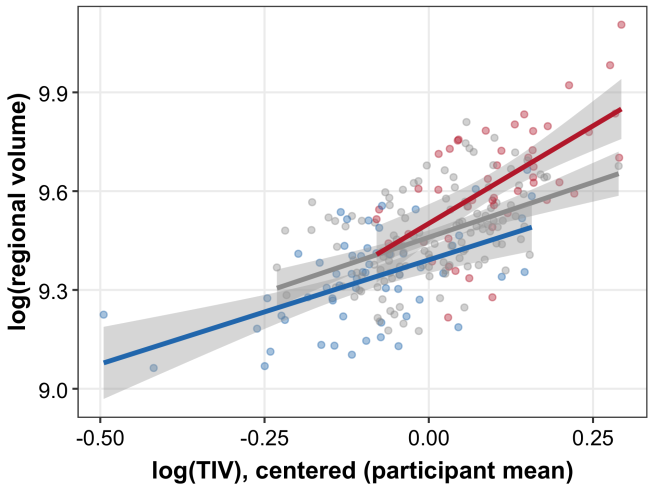


#### Right superior temporal

Deletion: Slope (slope p 0.003)
Duplication: Intercept (slope p 0.177)

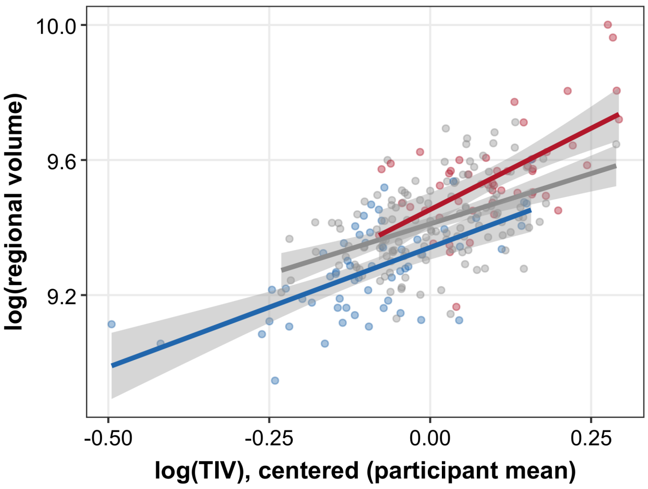


#### Left supramarginal

Deletion: n.s. screen
Duplication: n.s. screen

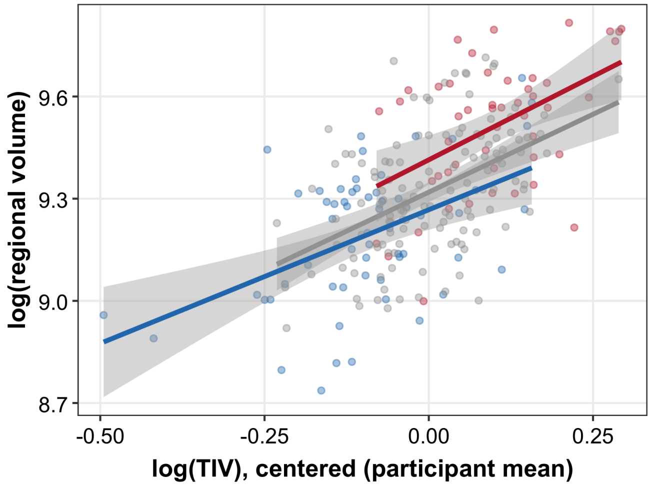


#### Right supramarginal

Deletion: n.s. screen
Duplication: n.s. screen

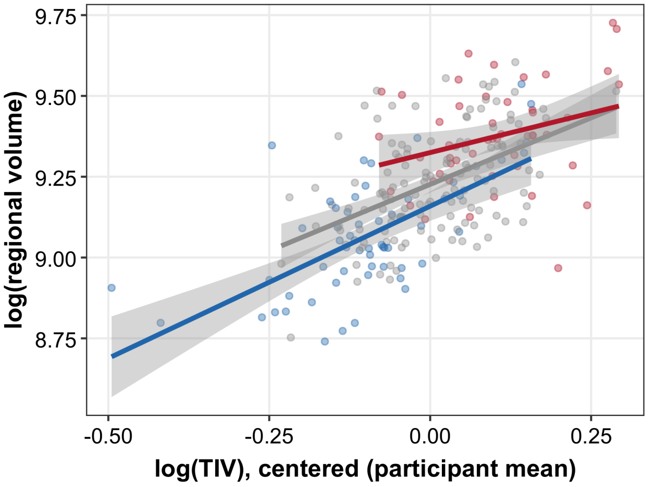


#### Left temporal pole

Deletion: n.s. screen
Duplication: n.s. screen

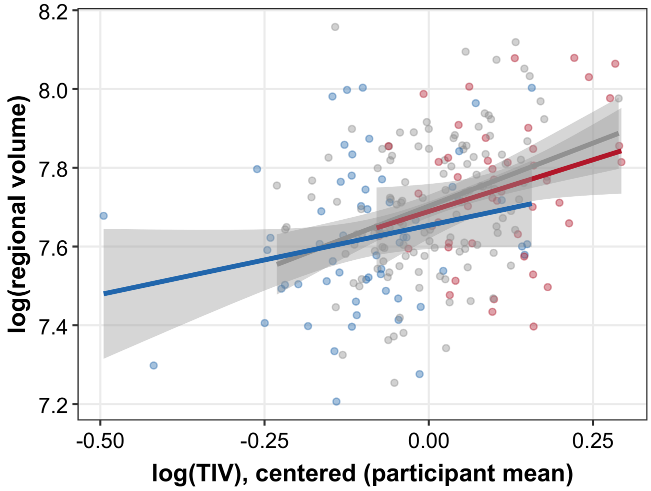


#### Right temporal pole

Deletion: n.s. screen
Duplication: n.s. screen

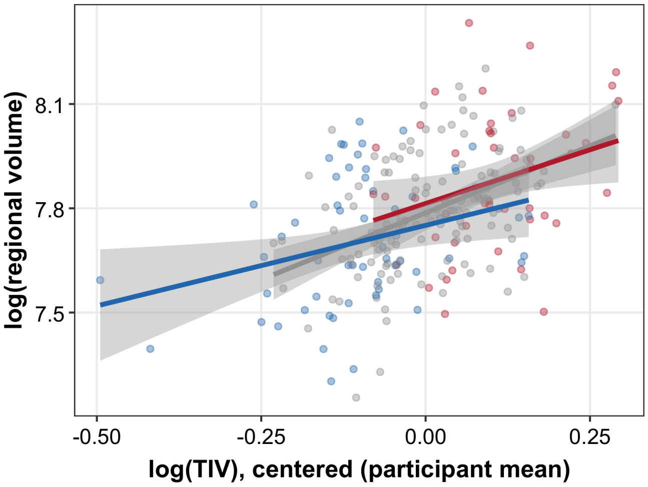

### Supplementary Figure S9. Regional allometric fits — cortical (12/12)

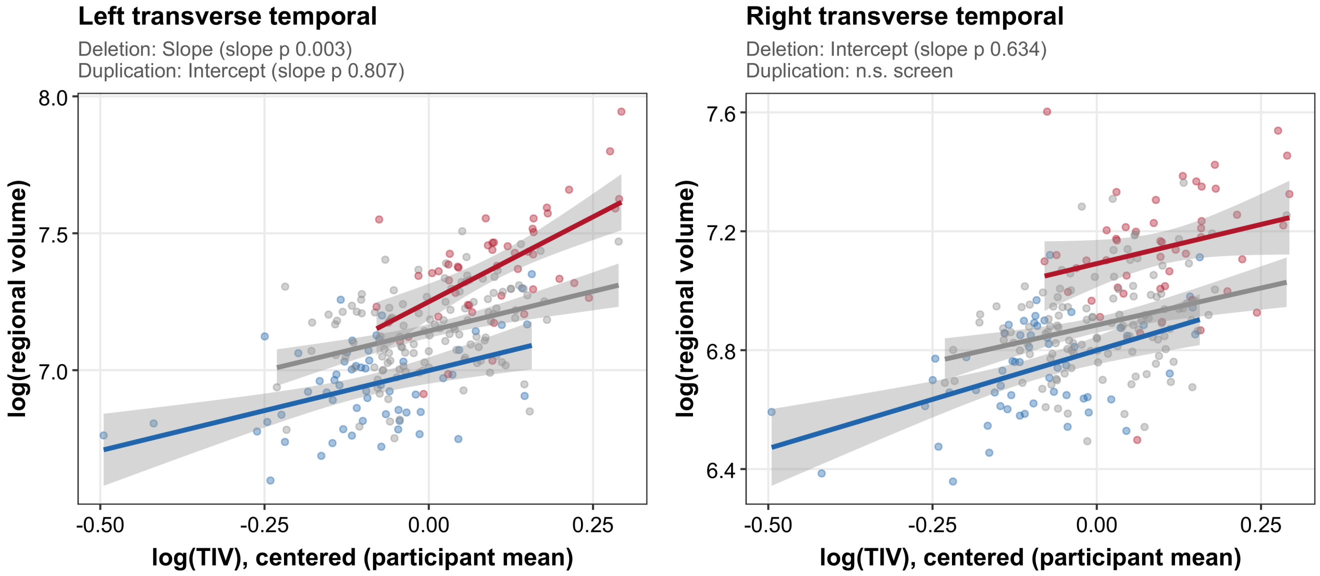

### Supplementary Figure S9. Regional allometric fits — subcortical (1/3)

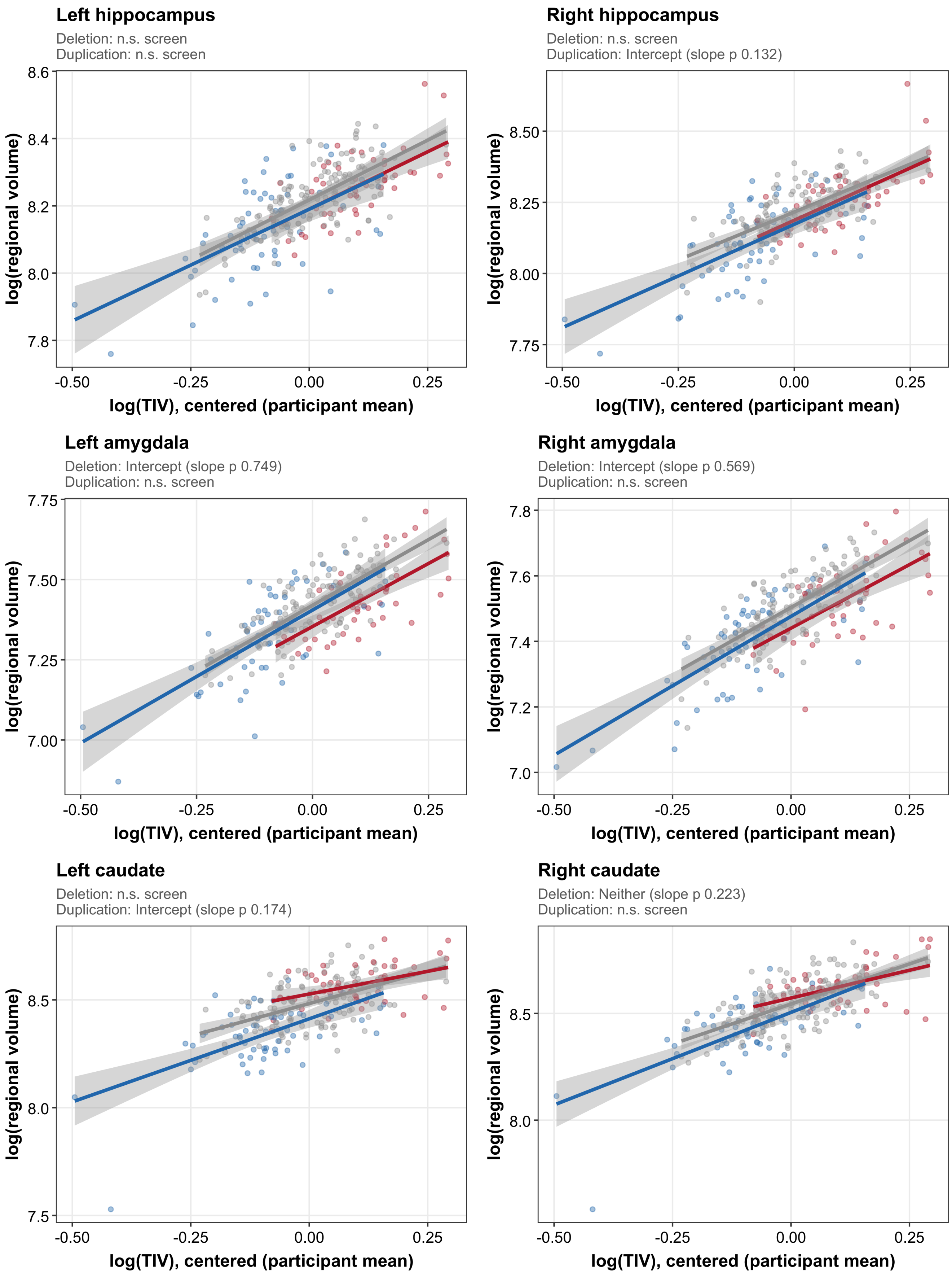

### Supplementary Figure S9. Regional allometric fits — subcortical (2/3)

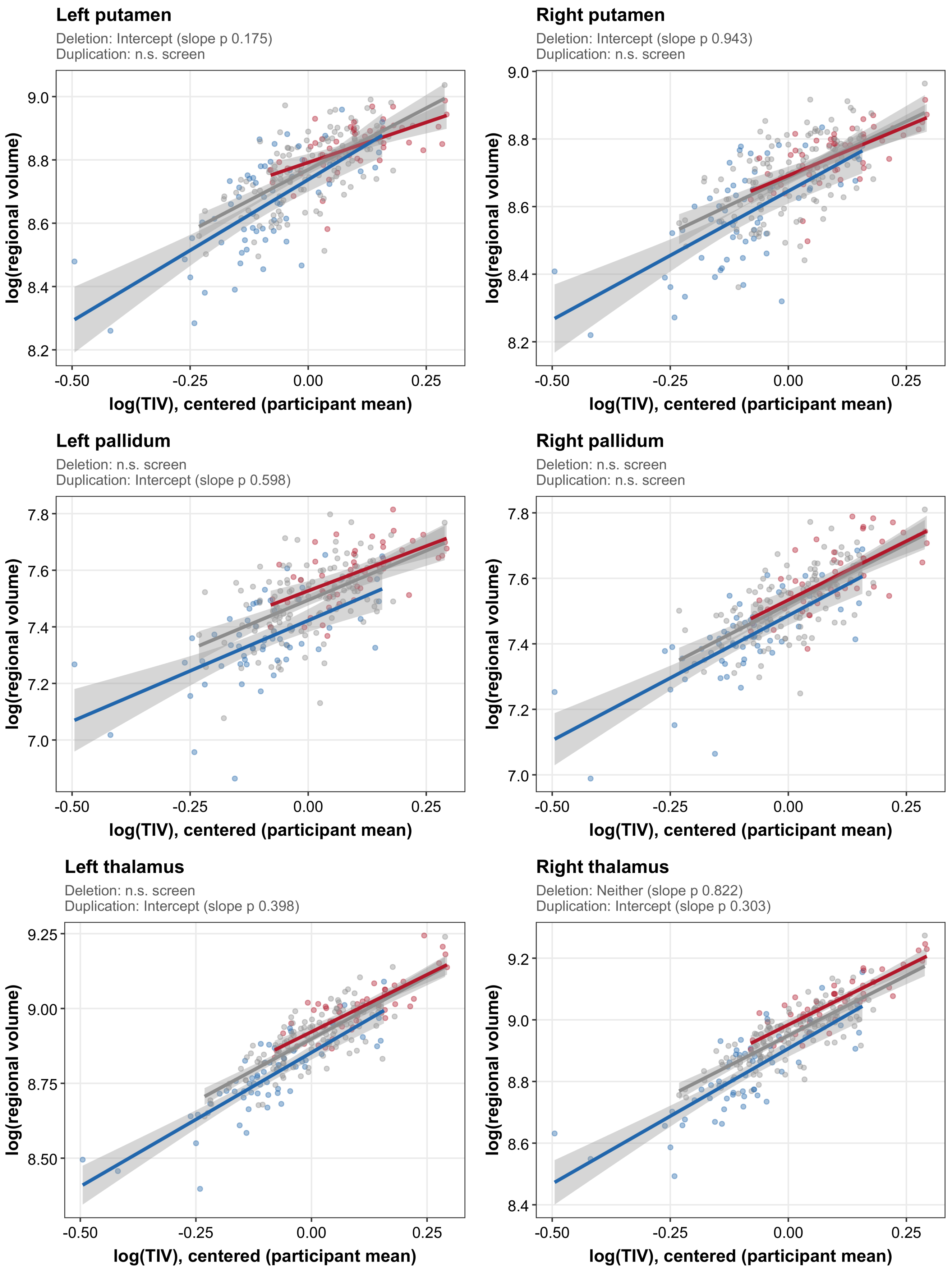

### Supplementary Figure S9. Regional allometric fits — subcortical (3/3)

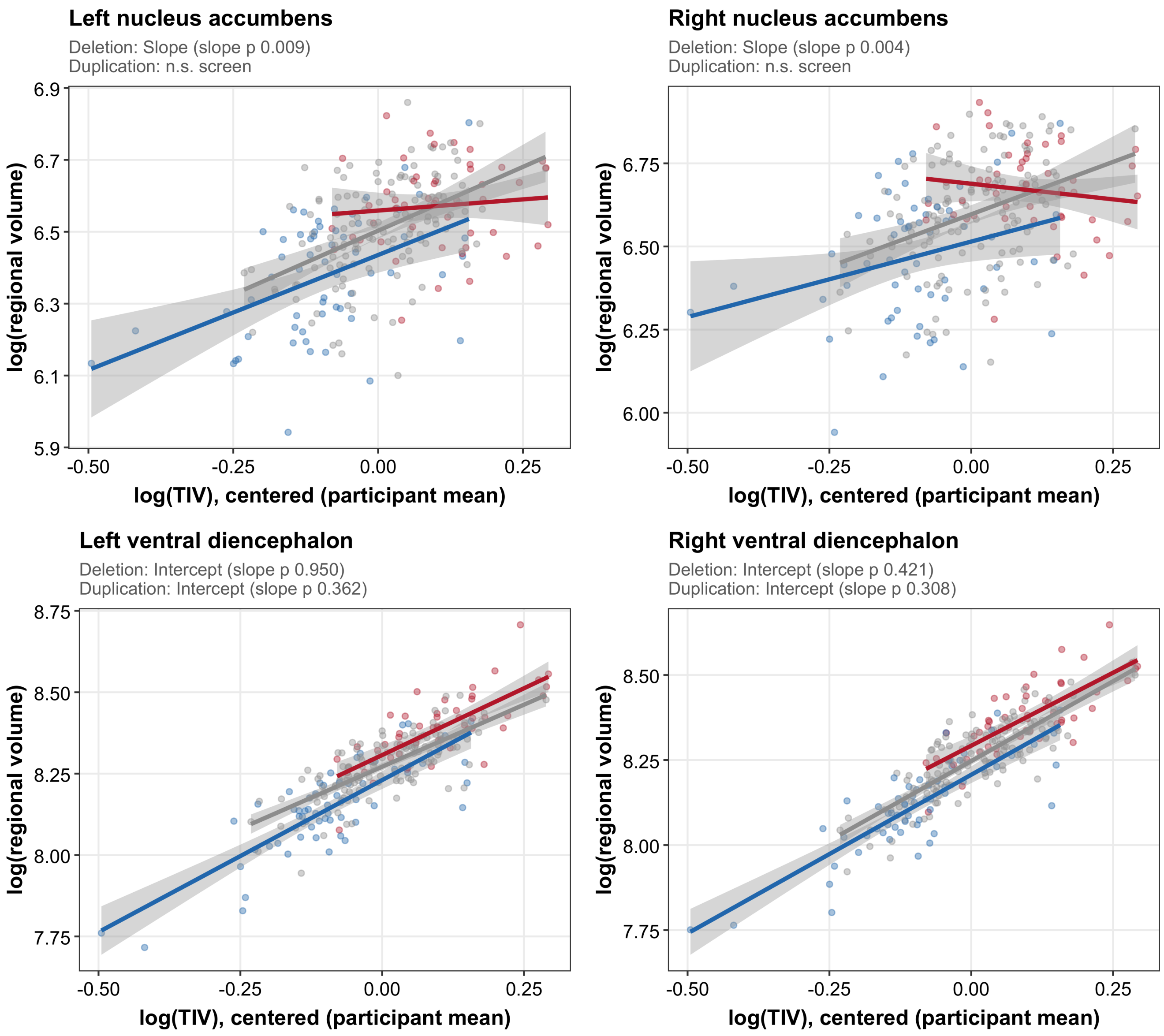

### Supplementary Figure S9. Regional allometric fits — global & other (1/4)

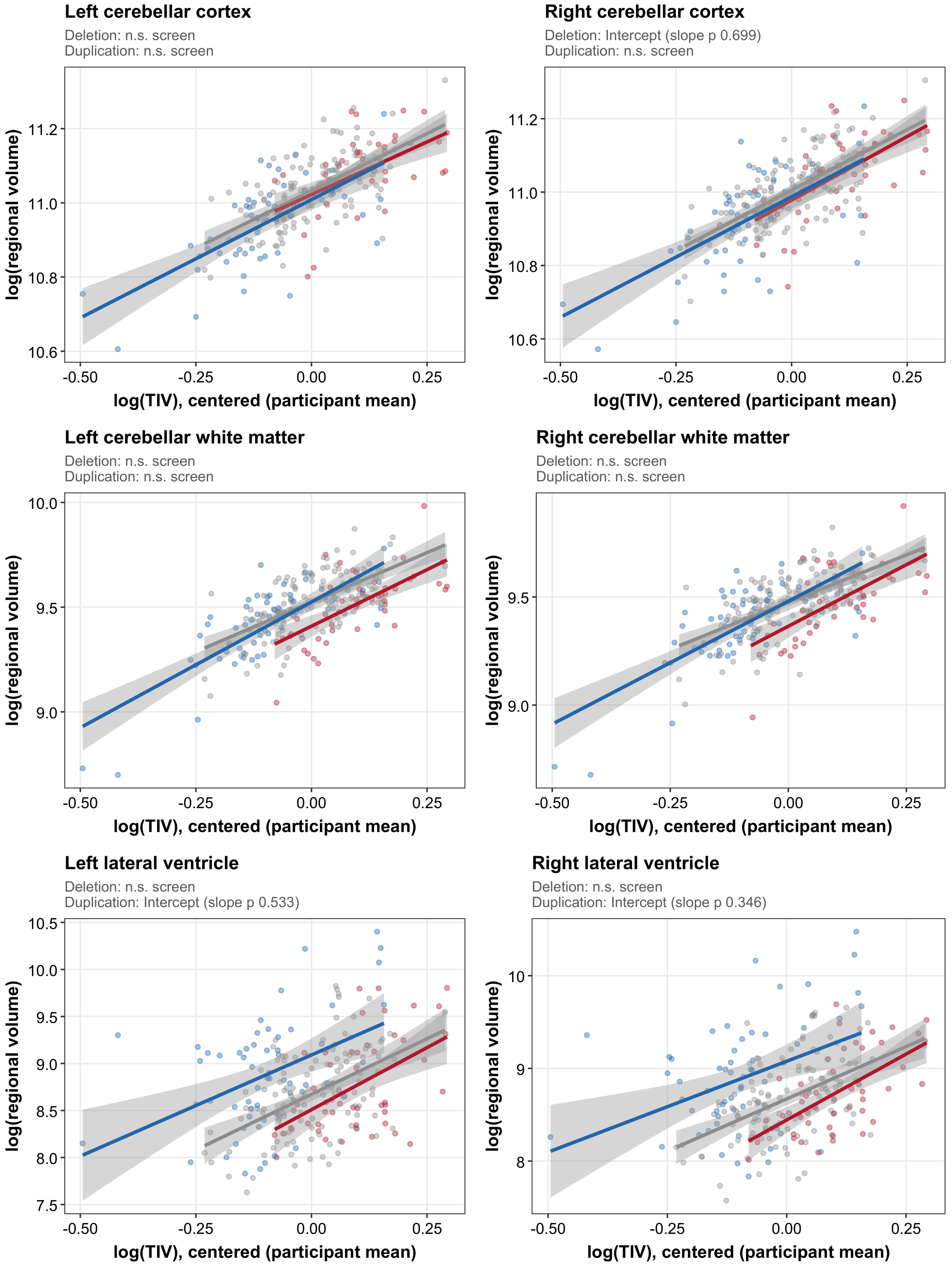

#### Supplementary Figure S9. Regional allometric fits — global & other (2/4)

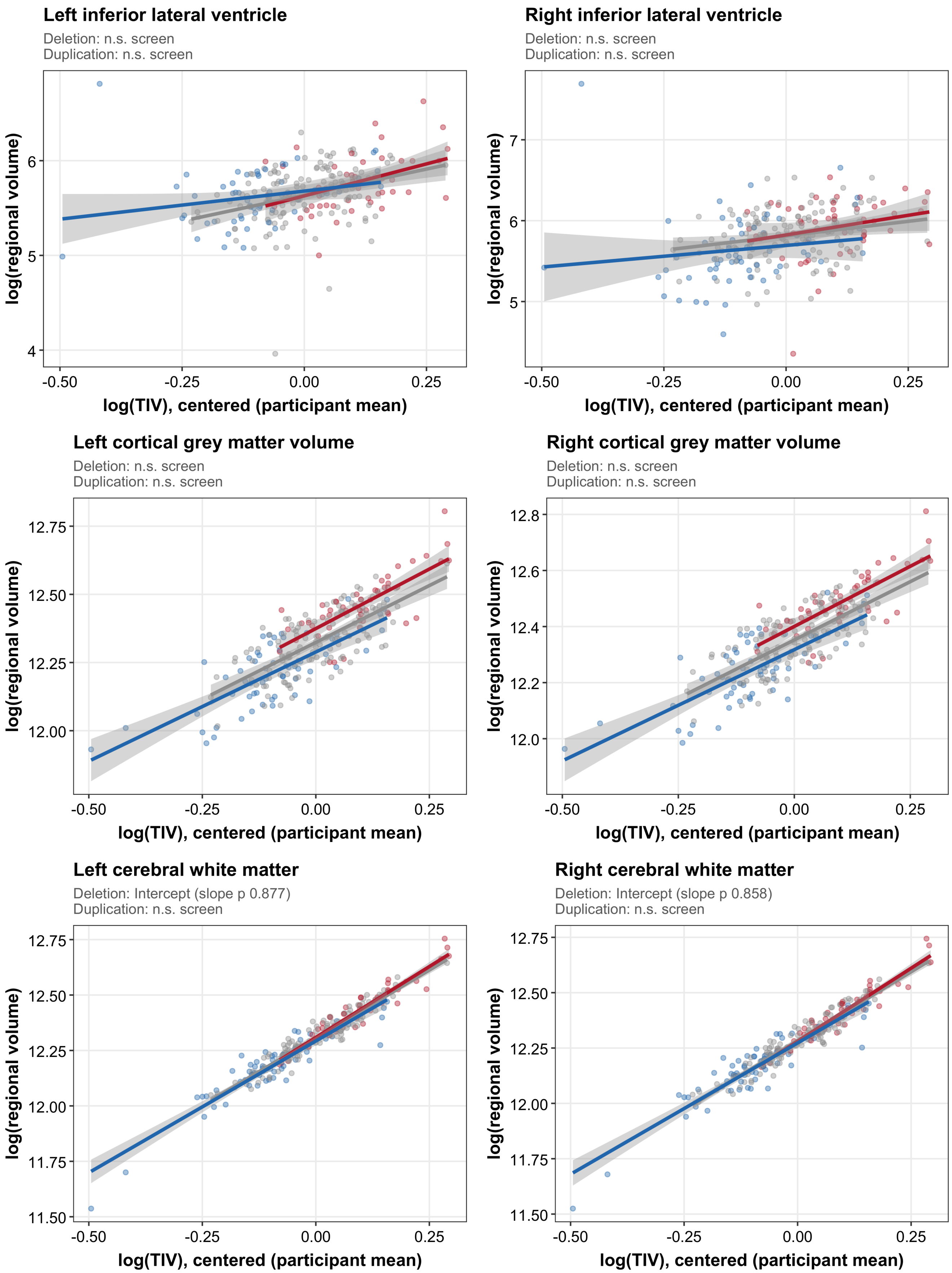

### Supplementary Figure S9. Regional allometric fits — global & other (3/4)

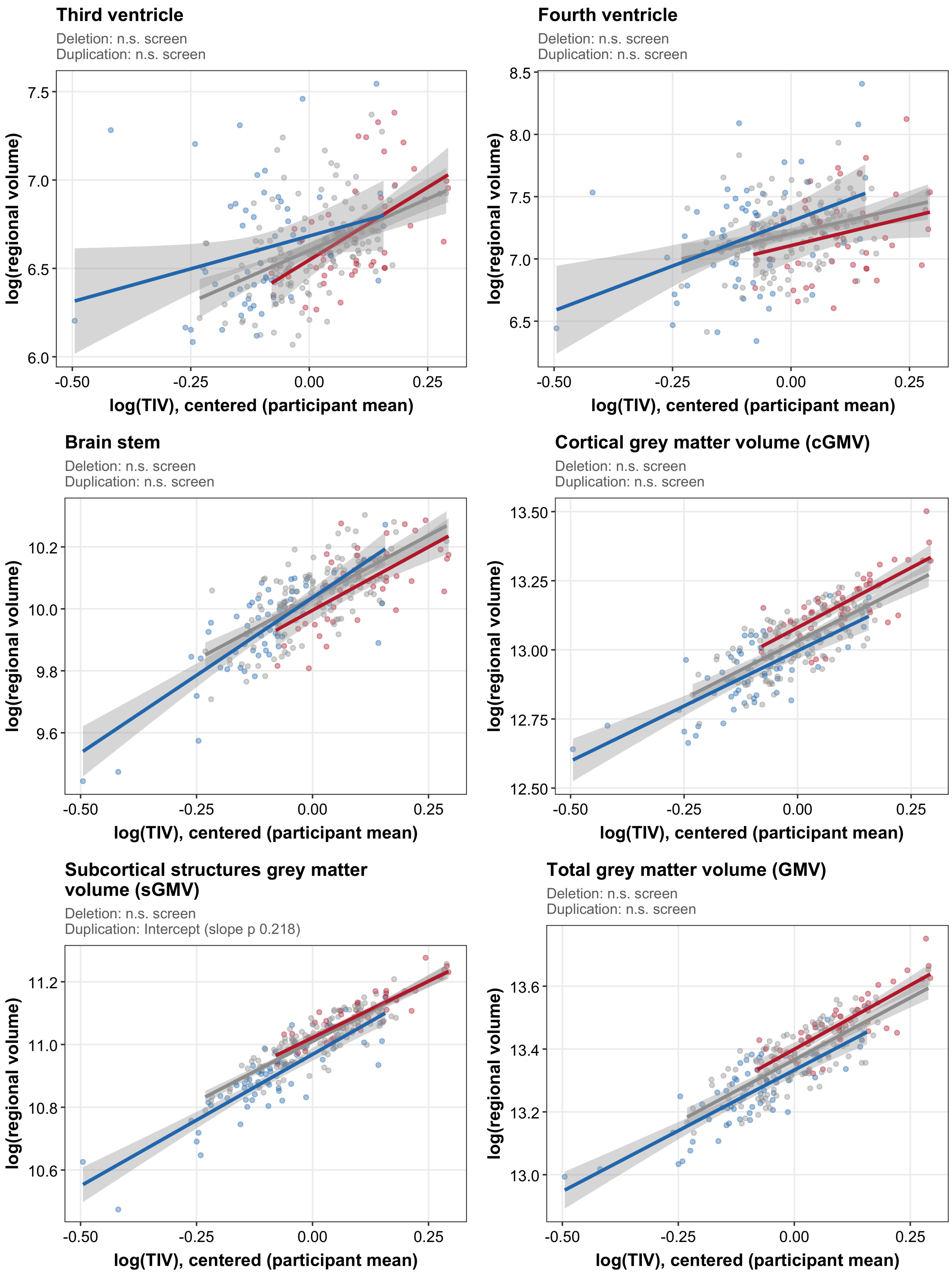

### Supplementary Figure S9. Regional allometric fits — global & other (4/4)

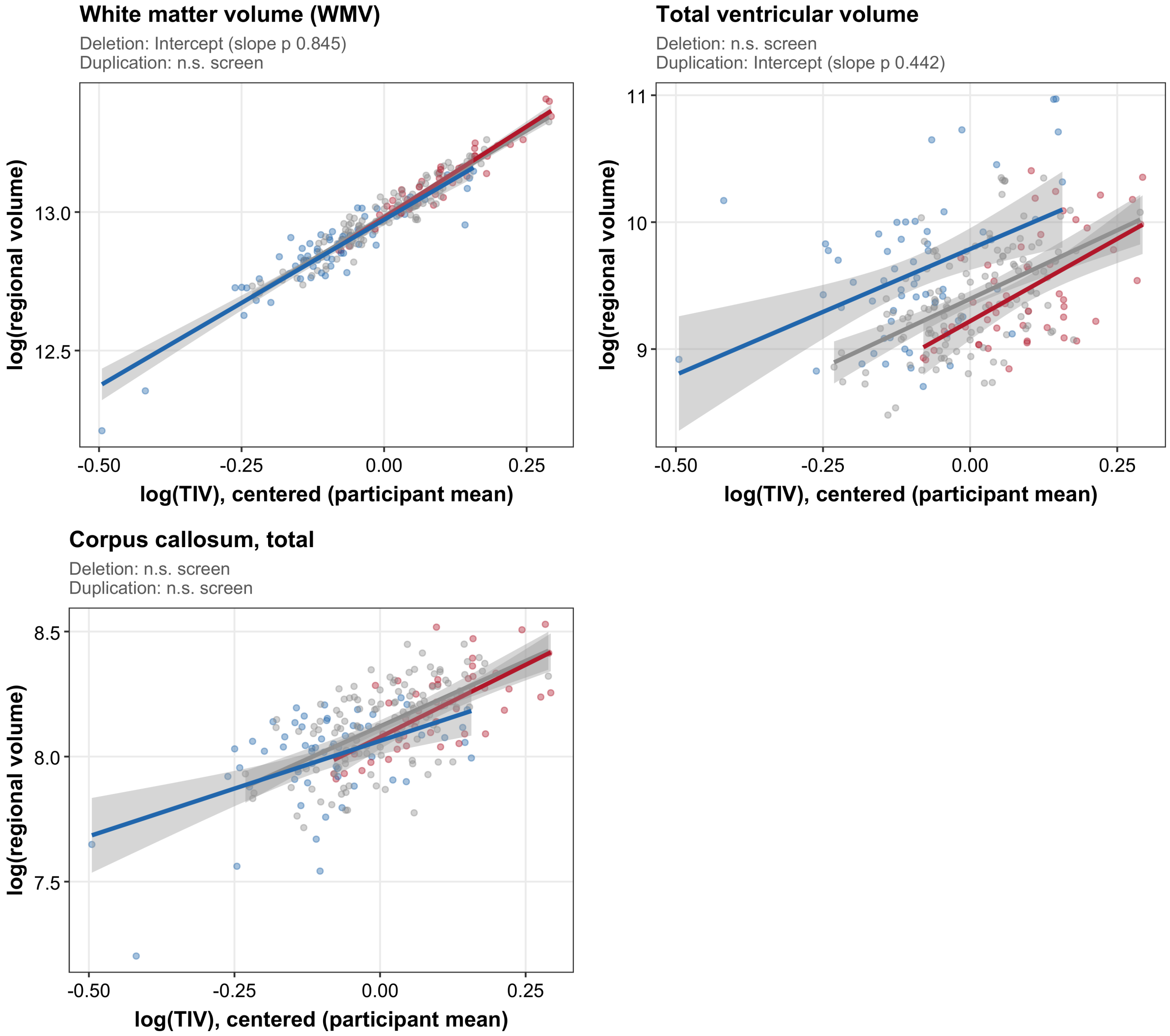


**Supplementary Figure S9: Regional visualization of allometric volume phenotypes**. Log-transformed regional volume is plotted against centered log(TIV) for controls (grey), deletion (red), and duplication (blue). Each point represents one participant; participants with two sessions are shown at their session mean. Lines show descriptive least-squares fits for each genotype with 95% confidence bands. Panel annotations report the allometric classification for each CNV: measures that did not pass the FDR-corrected CNV-specific omnibus screen are labeled n.s. screen; among screen-positive measures, those with a nominally significant genotype-by-size interaction are labeled slope, and those without an altered exponent but with a significant difference in the scaling constant are labeled intercept. The interaction p-value is shown for screen-positive measures.
